# Snx14 proximity labeling reveals a role in saturated fatty acid metabolism and ER homeostasis defective in SCAR20 disease

**DOI:** 10.1101/2020.05.31.126441

**Authors:** Sanchari Datta, Jade Bowerman, Hanaa Hariri, Rupali Ugrankar, Kaitlyn M. Eckert, Chase Corley, Gonçalo Vale, Jeffrey G. McDonald, Mike Henne

## Abstract

Fatty acids (FAs) are central cellular metabolites that contribute to lipid synthesis, and can be stored or harvested for metabolic energy. Dysregulation in FA processing and storage causes toxic FA accumulation or altered membrane compositions and contributes to metabolic and neurological disorders. Saturated lipids are particularly detrimental to cells, but how lipid saturation levels are maintained remains poorly understood. Here, we identify the cerebellar ataxia SCAR20-associated protein Snx14, an endoplasmic reticulum (ER)-lipid droplet (LD) tethering protein, as a novel factor required to maintain the lipid saturation balance of cell membranes. We show that *SNX14*^KO^ cells and SCAR20 disease patient-derived cells are hypersensitive to saturated FA (SFA)-mediated lipotoxic cell death that compromises ER integrity. Using APEX2-based proximity labeling, we reveal the protein composition of Snx14-associated ER-LD contacts and define a functional interaction between Snx14 and Δ-9 FA desaturase SCD1. Lipidomic profiling reveals that *SNX14*^KO^ cells increase membrane lipid saturation following exposure to palmitate, phenocopying cells with reduced SCD1 activity. In line with this, *SNX14*^KO^ cells manifest delayed FA processing and lipotoxicity, which can be rescued by SCD1 over-expression. Altogether these mechanistic insights reveal a role for Snx14 in FA and ER homeostasis, defects in which may underlie the neuropathology of SCAR20.

**Significance Statement:** SCAR20 disease is an autosomal recessive spinocerebellar ataxia primarily affecting children, and results from loss-of-function mutations in the *SNX14* gene. Snx14 is an endoplasmic reticulum (ER)-localized protein that localizes to ER-lipid droplet (LD) contacts and promotes LD biogenesis following exogenous FA treatment, but why Snx14 loss causes SCAR20 is unclear. Here, we demonstrate that following exposure to saturated fatty acids, Snx14-deficient cells have defective ER homeostasis and altered lipid saturation profiles. We reveal a functional interaction between Snx14 and fatty acid (FA) desaturase SCD1. Lipidomics shows Snx14-deficient cells contain elevated saturated lipids, closely mirroring SCD1-defective cells. Furthermore, SCD1 over-expression can rescue Snx14 loss. We propose that Snx14 maintains cellular lipid homeostasis, the loss of which underlies the cellular basis for SCAR20 disease.

## Introduction

Cells regularly internalize exogenous fatty acids (FAs), and must remodel their metabolic pathways to process and properly store FA loads. As a central cellular currency that can be stored, incorporated into membrane lipids, or harvested for energy, cells must balance FA uptake, processing, and oxidation to maintain homeostasis. Defects in any of these elevates intracellular free fatty acids which can act as detergents and damage organelles. Excessive membrane lipid saturation can also alter organelle function and contribute to cellular pathology, known as lipotoxicity [1, 2]. Failure to properly maintain lipid compositions and storage contributes to many metabolic disorders [3] including type 2 diabetes [4], obesity [5], cardiac failure [6, 7] and various neurological diseases [8].

Properties of FAs such as their degree of saturation and chain length are key determinants of their fate within the cell [9]. High concentrations of saturated FAs (SFAs) in particular are highly toxic, as their incorporation into organelles affects membrane fluidity and can trigger lipotoxicity and cell death [10-13]. To prevent this, cells desaturate SFAs into mono-unsaturated FAs (MUFAs) before they are subsequently incorporated into membrane glycerophospholipids or stored as triglycerides (TG) in lipid droplets (LDs). LD production provides a lipid reservoir to sequester otherwise toxic FAs, providing a metabolic buffer to maintain lipid homeostasis [14, 15].

As LDs are created by and emerge from the ER network, inter-organelle communication between the ER and LDs is vital for LD biogenesis [16]. Consequently numerous proteins that contribute to LD biogenesis, such as seipin [17, 18] and the diacylglyceride acyltransferase (DGAT) [19], are implicated in ER-LD crosstalk. Previously, we identified Snx14, a sorting nexin (SNX) protein linked to the cerebellar ataxia disease SCAR20 [20-22], as a novel factor that promotes FA-stimulated LD growth at ER-LD contacts [23, 24]. Snx14 is an ER-anchored integral membrane protein. During periods of elevated FA flux, Snx14 is recruited to ER-LD contact sites where it promotes the incorporation of FAs into TG as LDs grow [23]. In line with this, *SNX14*^KO^ cells exhibit defective LD morphology following oleate addition, implying Snx14 is required for proper FA storage in LDs. Related studies of Snx14 homologs in yeast and *Drosophila* indicate a conserved role for Snx14 family proteins in FA homeostasis and LD biogenesis [25, 26].

Despite these insights, why humans with Snx14 loss-of-function mutations develop the cerebellar ataxia disease SCAR20 remains enigmatic. Given the proposed role of Snx14 in lipid metabolism, and that numerous neurological pathologies arise through defects in ER lipid homeostasis [27-29], here we investigated whether Snx14 loss alters the ability of cells to maintain lipid homeostasis in response to FA influx. Our findings indicate that Snx14-deficient cells are hypersensitive to SFA exposure, and manifest defects in ER morphology and ER-associated lipid metabolism.

## Results

### *SNX14*^KO^ cells are hypersensitive to saturated fatty acid associated lipotoxicity

Previously, we showed that following oleate addition, Snx14 enriches at ER-LD contacts to promote LD growth, and Snx14 loss disrupts LD homeostasis [23]. To further dissect Snx14 function in maintaining lipid homeostasis, we interrogated how *SNX14*^KO^ cells respond to exposure to various saturated and unsaturated FAs. We exposed wildtype (WT) or *SNX14*^KO^ *U2OS* cells to titrations of specific SFAs or MUFAs for 48 hours, and monitored cell viability using an established crystal violet assay [30, 31]. Exposure to MUFAs including palmitoleate (16:1) and oleate (18:1) did not perturb cell viability of either WT or *SNX14*^KO^ cells even at 1000μM concentrations (**Fig 1 A, A’**). In contrast, treatment of WT cells with increasing concentrations of SFAs such as palmitate (16:0) or stearate (18:0) resulted in decreased cell survival, as previously reported [13, 32] (**Fig 1 B, B’**).

**Figure 1:**
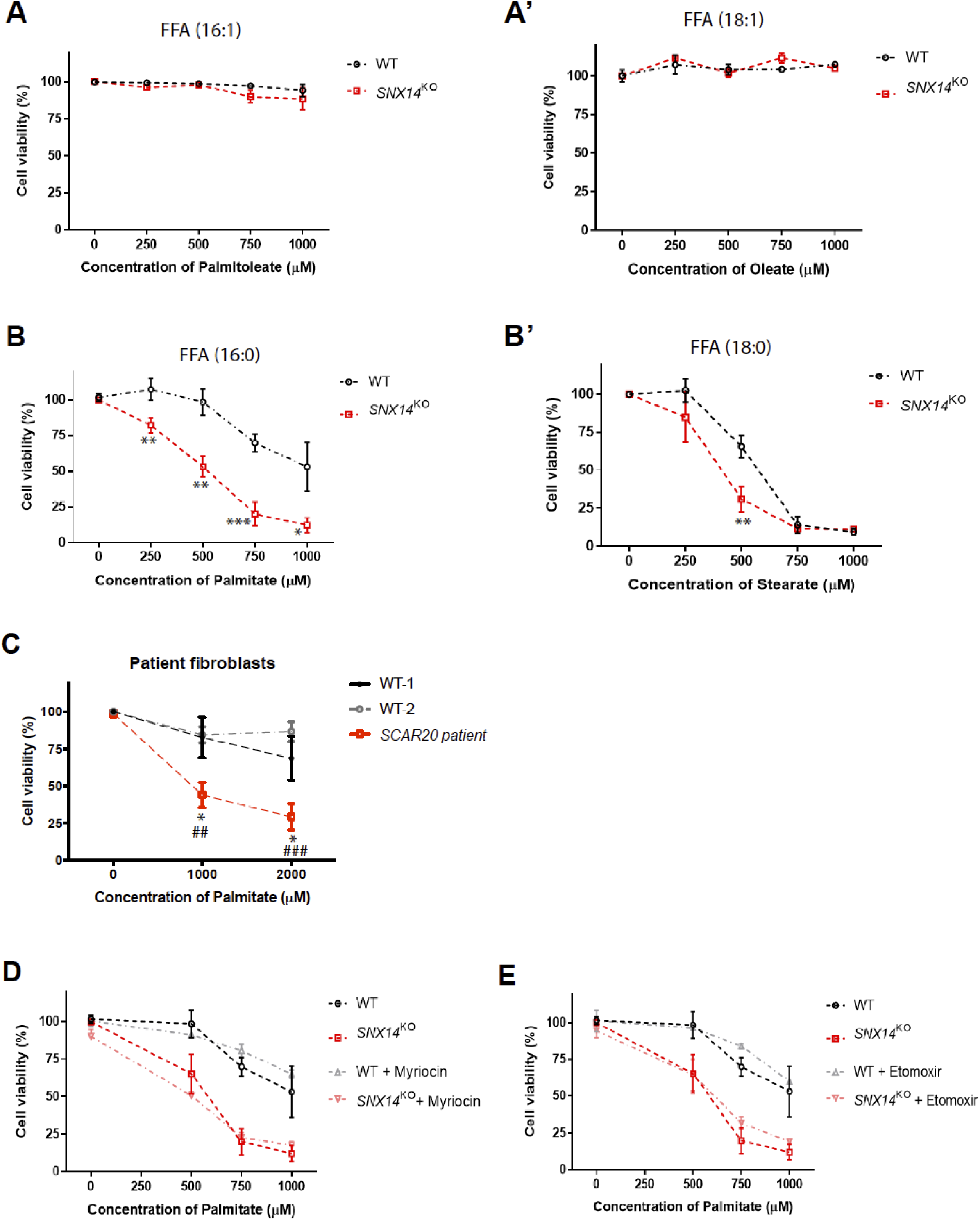
Snx14 deficient cells are hypersensitive to saturated fatty acids. A. Percentage of surviving cells denoted as cell viability (%) of WT and *SNX14*^KO^ cells, following treatment with increasing concentration (0, 250, 500, 750, 1000μM) of palmitoleate (FA|16:1) for 2 days. A’: Treatment of WT and *SNX14*^KO^ cells with similar increasing concentration of oleate (FA|18:1) for 2 days. The assay was repeated thrice in triplicates. Values represent mean±SEM. B. Cell viability (%) of WT and *SNX14*^KO^ cells, showing *SNX14*^KO^ cells are hypersensitive following addition of increasing concentration (0, 250, 500, 750, 1000μM) of palmitate (FA|16:0) for 2 days. B’: Exposure to similar increasing concentration of stearate (FA|18:0) for 2 days in WT and *SNX14*^KO^ cells. The assay was repeated thrice in triplicates. Values represent mean±SEM. Significance test between WT and *SNX14*^KO^ (n=3, ***p<0.0001,**p<0.001,*p<0.01, multiple t-test by Holm-Sidak method with alpha = 0.05) C. Cell viability (%) of fibroblasts derived from 2 WT (*SNX14*^+^/*SNX14*^+^) subjects and 1 SCAR20 patient with homozygous recessive mutations in *SNX14*, showing hypersensitivity of SCAR20 cells following 2-day exposure to 0, 1000, 2000μM palmitate. The assay was repeated thrice in triplicates. Values represent mean±SEM (n=3, *p<0.01 relative to WT-1; ^##^p<0.001 and ^###^p<0.0001 relative to WT-2; multiple t-test by Holm-Sidak method with alpha = 0.05). D. Cell viability (%) of WT and *SNX14*^KO^ cells following exposure to increasing palmitate concentration (0, 250, 500, 750, 1000 μM) and treated with 10 μM Etomoxir for 2 days. The assay was repeated thrice in triplicates. Values represent mean±SEM. E. Cell viability (%) of WT and *SNX14*^KO^ cells following addition of increasing palmitate concentration (0, 250, 500, 750, 1000μM) in presence of 50μM Myriocin for 2 days. The assay was repeated thrice in triplicates. Values represent mean±SEM.

Intriguingly, *SNX14*^KO^ *U2OS* cells were hyper-sensitive to SFA-induced death compared to WT cells (**Fig 1 B, B’**). The concentration of palmitate at which ∼50% of WT cells survive is ∼1000μM, but only ∼500μM for *SNX14*^KO^ (**Fig 1B**). Similarly, exposure to ∼600μM stearate resulted in ∼50% cell viability for WT cells, but only required ∼300μM for *SNX14*^KO^ (**Fig 1B’**). Consistent with this, SCAR20 patient-derived fibroblasts [20, 24] which are homozygous for loss-of-function Snx14 mutations exhibited significantly reduced cell viability compared to control fibroblasts following palmitate addition (**Fig 1C**). SCAR20 patient cells exhibited ∼50% viability at ∼1000μM palmitate exposure, whereas more than 50% WT cells were viable even at 2000μM palmitate treatment. Collectively, these observations indicate that Snx14-deficient cells are hyper-sensitive to SFA exposure.

Once internalized, free FAs (FFAs) are esterified and shunted into several distinct metabolic fates, including their incorporation into ceramides [33], glycerophospholipids [34], or neutral lipids[35]. They can also be harvested by catabolic breakdown in oxidative organelles like mitochondria [2]. To begin to dissect why *SNX14*^KO^ cells were hyper-sensitive to SFAs, we conducted a systemic analysis of each FA-associated pathway. Since intracellular ceramide accumulation can itself be toxic [33], we tested whether pharmacologically lowering ceramide synthesis could rescue *SNX14*^KO^ SFA-associated toxicity. We treated cells with myriocin, which inhibits the SPT complex that incorporates palmitate into newly synthesized ceramides [10]. Myriocin treatment did not rescue *SNX14*^KO^ SFA hypersensitivity, suggesting that elevated ceramides does not contribute to *SNX14*^KO^ palmitate-induced cell death (**Fig 1D**). Next, we examined whether mitochondrial FA oxidation was required for *SNX14*^KO^ hypersensitivity by treating cells with etomoxir that inhibits mitochondrial FA uptake [36, 37]. This did not suppress palmitate-induced lipotoxicity in *SNX14*^KO^ cells, indicating perturbed mitochondrial FA oxidation is likely not causative of Snx14-associated lipotoxicity (**Fig 1E**).

### SFA-induced lipotoxicity in *SNX14*^KO^ cells is associated with defects in ER morphology

A major FA destination is their incorporation into membrane glycerophospholipids via *de novo* lipid synthesis at the ER. Excessive SFA incorporation into diacyl-glycerophospholipids can impact ER membrane fluidity and drive cellular stress [12, 38]. To determine whether SFA exposure affected ER homeostasis in *SNX14*^KO^ cells, we examined ER morphology of WT and *SNX14*^KO^ cells exposed to palmitate via fluorescence confocal microscopy. Whereas the immunofluorescently stained ER network of WT cells was reticulated and regular following overnight exposure to 500μM palmitate, *SNX14*^KO^ cells displayed perturbed ER morphology. The ER of *SNX14*^KO^ cells appeared fragmented, and contained drastic bulges within the tubular network following palmitate treatment (**Fig 2A; red arrows**). A subpopulation of *SNX14*^KO^ cells even exhibited solubilized ER lumen marker in the cytoplasm, suggesting drastic defects in ER integrity. We quantified these ER morphological alterations into distinct classes (**Fig 2A**,**B; white arrows indicating ER morphology of each class**), revealing ∼70% of *SNX14*^KO^ cells exhibited irregular or fragmented ER structure following palmitate exposure, compared to only ∼22% of WT (**Fig 2B**). These perturbations became more prominent when examining ER ultrastructure with transmission electron microscopy (TEM). Here, WT and *SNX14*^KO^ cells were either left untreated or cultured in media containing 500μM of palmitate for 6hrs or 16hrs. Even with palmitate, WT cells exhibited normal tubular ER networks and did not manifest any significant change in the ER morphology (**Fig 2C**). *SNX14*^KO^ cells not exposed to palmitate also exhibited normal ER morphology. However, the ER of *SNX14*^KO^ cells exposed to 6hrs palmitate appeared swollen and dilated, forming sheet-like extensions within the thin-section plain (**Fig 2C; red arrows**). This ER dilation was more pronounced following 16hrs treatment (**Fig 2C; red arrows**).

**Figure 2:**
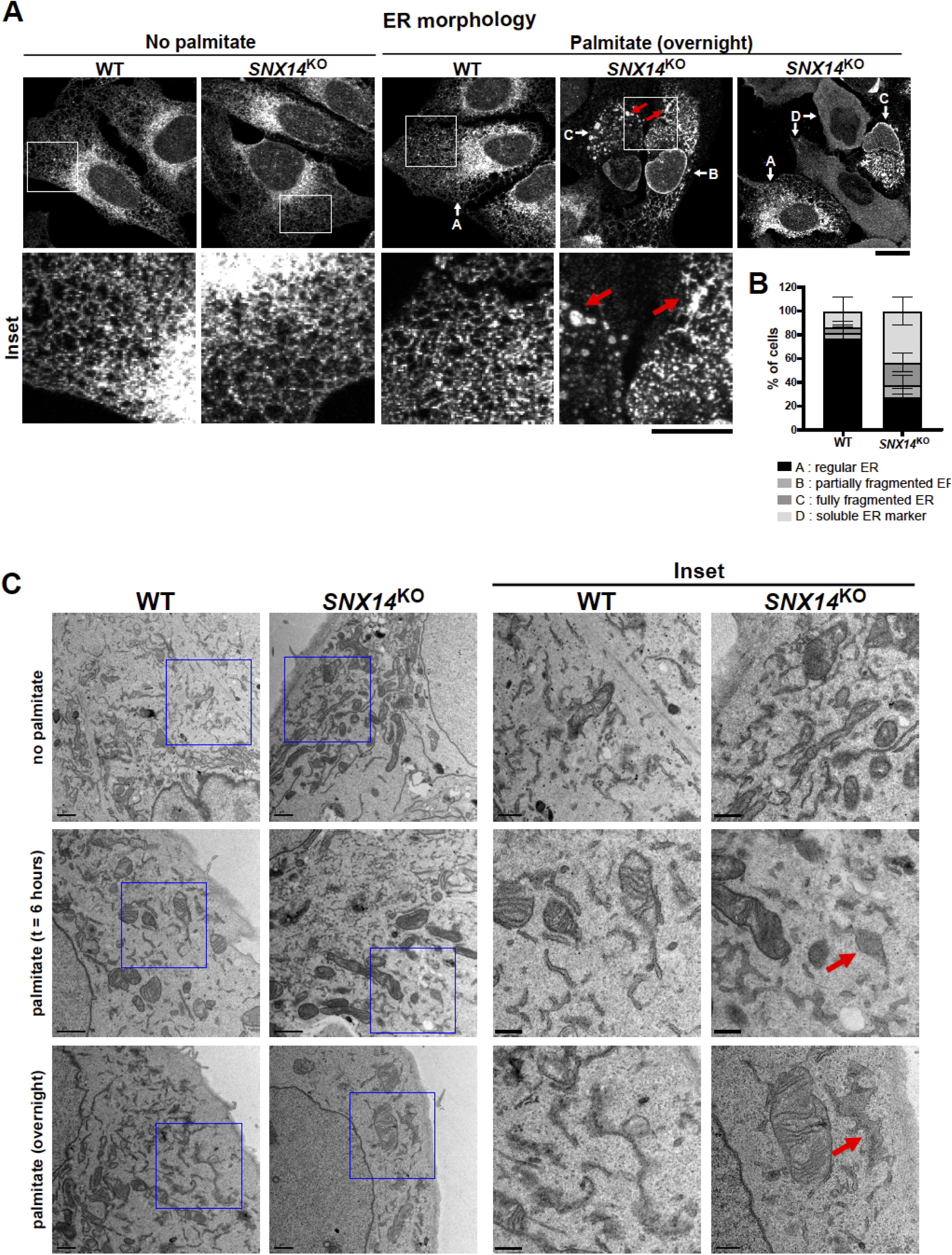
Palmitate-induced hypersensitivity in *SNX14*^KO^ is associated with defective ER morphology. **A**. Immunofluorescent (IF) labeling of the ER with α-HSP90B1 (ER marker) antibody before and after overnight palmitate treatment in WT and *SNX14*^KO^ cells. Scale bar = 10μm. Red arrows indicating the drastic bulges in the ER. White arrows indicating each class of ER morphology where A is regular ER, B is partially fragmented ER, C is fully fragmented ER and D shows soluble ER marker. **B**. Percentage of palmitate treated WT and *SNX14*^KO^ cells quantified and grouped based on whether the ER morphology is regular (A), partially fragmented (B), fully fragmented (C) or completely soluble (D). Total ∼100 cells quantified from 3 experiments. Values represent mean±SEM. **C**. TEM micrographs of WT and *SNX14*^KO^ cells with no palmitate treatment and with palmitate treatment for 6hr and overnight to visualize ER ultra-structure. Scale bar = 1μm. Scale bar of inset = 0.5 μm. Red arrows indicate ER dilation in palmitate treated *SNX14*^KO^ cells.

Since Snx14 is implicated in LD biogenesis, we also examined LD morphology in WT and *SNX14*^KO^ cells exposed to palmitate. Whereas WT cells generated small LDs in response to palmitate, *SNX14*^KO^ cells exhibited significantly fewer LDs that stained poorly with the LD dye monodansylpentane (MDH), suggesting defective palmitate processing and LD incorporation (**Fig S2A, B**). Collectively, these observations suggest that *SNX14*^KO^ cells manifest altered ER architecture and LD homeostasis following prolonged exposure to palmitate.

Changes in ER lipid homeostasis can induce the unfolded protein response (UPR) as well as caspase-dependent apoptotic cell death [13]. To understand whether such responses were associated with palmitate-induced cell death in Snx14 deficienct cells, we monitored them in *SNX14*^KO^ cells. First, we examined the levels of spliced Xbp1 (s-Xbp1) following palmitate exposure. As expected, palmitate elevated s-Xbp1 levels compared to no treatment in both WT and *SNX14*^KO^ cells, but there was not a significant difference between the two cell lines (**Fig S2C**). In line with this, addition of the IRE1 inhibitor 4μ8c or PERK inhibitor GSK2606414, both which can suppress branches of UPR signaling, did not rescue palmitate-induced cell death in *SNX14*^KO^ cells, suggesting altered or hyper-active UPR signaling was not causative of *SNX14*^KO^ cell death following palmitate exposure (**Fig S2D**). To dissect whether SFAs induced an apoptotic response in *SNX14*^KO^ cells, we also treated cells with the caspase-6/8 inhibitor SCP0094. This did not rescue palmitate-induced cell death, indicating *SNX14*^KO^ cells were not manifesting hyper-active caspase-6/8-dependent apoptosis (**Fig S2D**).

### APEX-based proteomics reveals Snx14 is in proximity to proteins involved in SFA metabolism

Given that Snx14 was required for maintaining ER morphology and LD biogenesis following palmitate exposure, we next investigated what proteins Snx14 interacted with that may promote lipid homeostasis. Previously we utilized a proximity-based technology based on the ascorbate peroxidase APEX2 to examine the localization of APEX2-tagged Snx14 at ER-LD contact sites using TEM [23]. APEX2-tagging also enables the local interactome of a protein-of-interest to be interrogated. The addition of biotin-phenol and hydrogen peroxide to APEX2-expressing cells induces the local biotinylation of proteins within ∼20nm of the APEX2 tag. These biotinylated proteins can be subsequently affinity purified via streptavidin beads and identified via mass-spectrometry (MS) [39] (**Fig 3A**). As ER-LD contacts are lipogenic ER sub-domains with known roles in FA processing, we hypothesized that Snx14’s enrichment at ER-LD junctions represented an opportunity to use APEX2 technology to reveal proteins that contribute to ER lipid metabolism in conjunction with Snx14.

**Figure 3:**
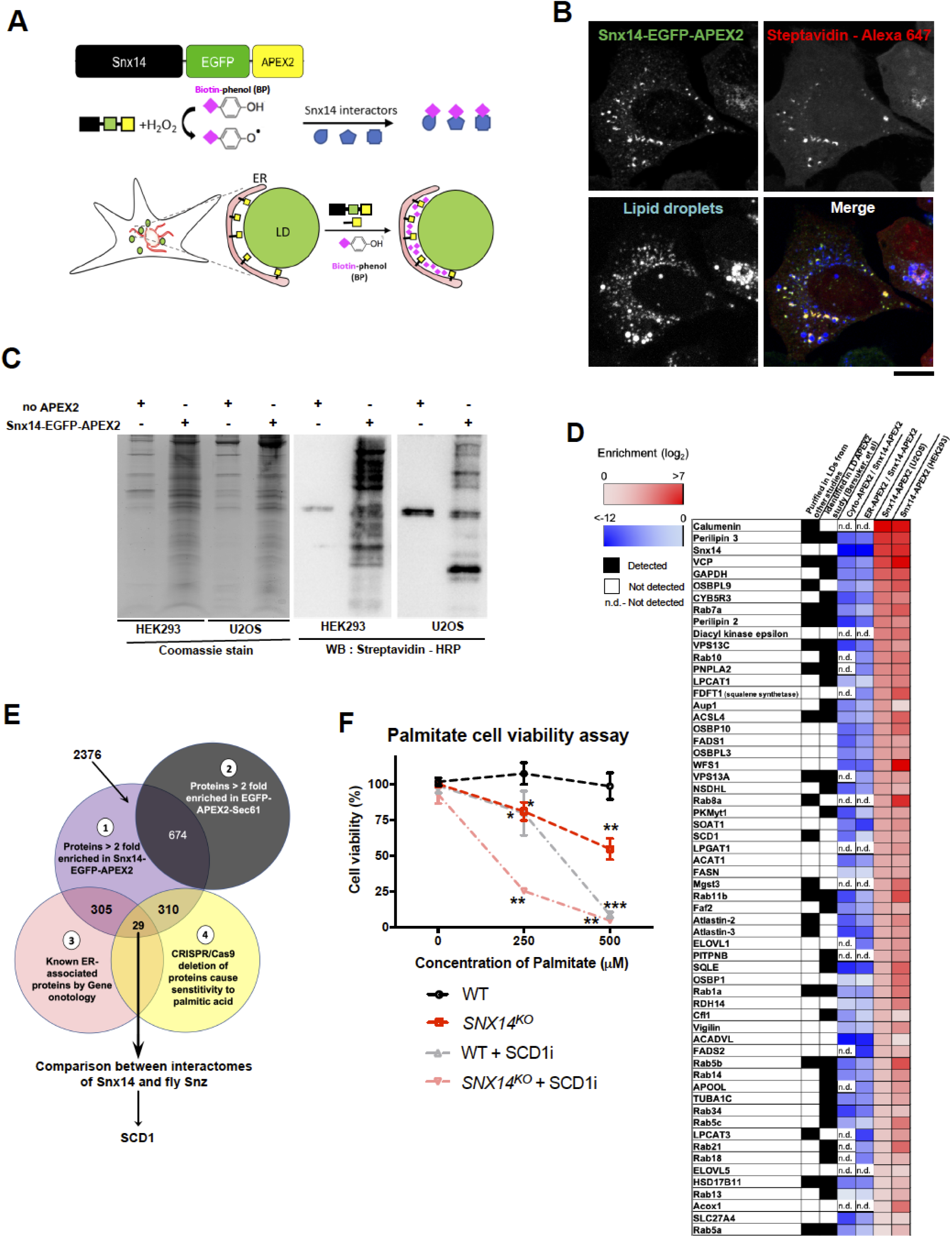
APEX2-based proteomics reveals the Snx14-associated ER-LD proteome. **A**. Schematic diagram showing Snx14 tagged with EGFP followed by APEX2 at the C-terminus. The reaction of biotin phenol and H_2_O_2_ catalysed by APEX2 generates biotin-phenoxyl radicals which covalently attaches with proteins in proximity of ∼20nm from Snx14. Following oleate treatment, APEX2 tagged Snx14 enriches at ER-LD contacts and hence the labelling reaction is expected to biotinylate the interactors of Snx14 at those contacts. **B**. Co-IF staining of cells stably expressing Snx14-EGFP-APEX2 with anti-EGFP (green) antibody, streptavidin-conjugated Alexa647 fluorophore (biotinylated proteins, red) antibody and LDs stained with monodansylpentane (MDH, blue) followed by confocal imaging revealed colocalization of Snx14 and biotinylated proteins surrounding LDs. Scale bar = 10μm. **C**. Biotinylated proteins pulled down by streptavidin conjugated beads from HEK293 and U2OS cells expressing Snx14-EGFP-APEX2 and no-APEX2 (negative control) were coomassie stained. Western blotting of the same pulled down lysates with streptavidin-HRP antibody revealed biotinylation of several proteins in Snx14-EGFP-APEX2 relative to the negative control. **D**. Heat map of ∼60 proteins highly enriched in Snx14-APEX2 proteomics and cross referenced with other LD proteomics study [40-44] (black box represents detected and white box represents no detection). The heatmap also shows the enrichment of the proteins in Snx14-APEX2 proteomics (red columns) as well as the de-enrichment in the cyto-APEX2 and ER-APEX2 proteomics (blue columns) (n.d. is not detected). **E**. Venn diagram representing the multi-stage analysis of the Mass spectrometry data which identified protein abundance in Snx14-EGFP-APEX2, no-APEX2 and EGFP-APEX2-Sec61. 2376 proteins are enriched >2 fold in Snx14-EGFP-APEX2 (1: purple circle). Among them 674 which are also enriched in APEX2 tagged Sec61 are excluded (2 : black circle). 305 of (1) are ER-associated according to Gene ontology analysis (3: red circle). Again 310 of (1) comprise of genes whose CRISPR/Cas9 deletion cause palmitate sensitivity [46] (4 : yellow circle). Overlap of sets (1),(3) and (4) consist of 29 ER-associated proteins which are enriched for Snx14 and important for palmitate metabolism. This set belong to Snx14 interactome and is compared with its fly homolog Snz interactome. This final comparison indicated SCD1 as one of the top candidates. **F**. Cell viability of WT and *SNX14*^KO^ cells treated with 4μM SCDi along with increasing palmitate concentration (0, 250, 500μM) for 2 days. D’. The assay was repeated thrice in triplicates. Values represent mean±SEM. (n=3, *p<0.1, **p<0.001, ***p<0.0001, multiple t-test by Holm-Sidak method with alpha = 0.05)

We generated *U2OS* cells stably expressing Snx14-EGFP-APEX2 and *HEK293* cells transiently expressing Snx14-EGFP-APEX2 and exposed them to oleate to induce Snx14 recruitment to ER-LD contacts. We then treated cells with biotin-phenol for 30 minutes and hydrogen peroxide for 1 minute. As expected, co-immunofluorescence staining of *U2OS* cells for Streptavidin-Alexa647 and EGFP revealed their colocalization, suggesting proteins in close proximity to Snx14-EGFP-APEX2 were biotinylated (**Fig 3B**). Biotinylated proteins from both *U2OS* and *HEK293* cells were then affinity purified with streptavidin beads. Gel electrophoresis followed by Coomassie staining and anti-streptavidin/HRP Western blotting revealed many biotinylated proteins in Snx14-EGFP-APEX2-expressing cell lysates, but not in controls lacking APEX2 (**Fig 3C**). The bead enriched biotinylated proteins from both Snx14-EGFP-APEX2 and the control lacking APEX2 were then identified by tandem MS/MS proteomic analysis.

Previous proteomics studies have isolated LDs and characterized LD-associated proteins [40-44], and several of these were also detected in our APEX2-based approach (**Fig 3D**). However, to confirm that these peptide hits were specific to the Snx14-EGFP-APEX2 interactome, we also conducted proteomics on biotinylated proteins from cells expressing a soluble EGFP-APEX2 (cyto-APEX) as well as cells expressing APEX2-tagged Sec61β, a general ER marker (ER-APEX). High abundance peptides in the Snx14-EGFP-APEX2 proteomics that were correspondingly low in the cyto-APEX and ER-APEX were thus considered high confidence hits (**Fig 3D, Supplementary Table 1**). Notably, this list included well-characterized LD surface proteins Perilipin 3 (PLIN3), Perilipin 2, and PNPLA2. In fact, PLIN3 was one of the most enriched proteins from both cell line samples (>65-fold) (SFig 3A), consistent with it being a highly abundant LD protein that coats newly synthesized LDs. Proteins recently highlighted as localizing to ER-LD contact sites were also identified including VPS13C and VPS13A [45]. We also detected several Rab proteins such as Rab7a, Rab10, and Rab11b, which had previously been detected on LDs using a LD-targeted APEX approach [44]. Additionally, we detected enzymes involved in ER-associated lipid synthesis, including lyso-phospholipid acyltransferases LPCAT1, LPCAT3, and LPGAT, as well as enzymes driving sterol metabolism like the sterol O-acyltransferase SOAT1, the squalene synthase FDFT1, and the squalene epoxidase SQLE. Lastly, we noted several enzymes involved in FA metabolism and desaturation including ACSL4, ELOVL1, ELOVL5, FASN, SCD1, FADS1, and FADS2.

To identify Snx14 functional interactors that may play a role in FA metabolism that was defective in *SNX14*^KO^ cells, we implemented a multi-stage analysis approach (**Fig 3E**). First, we selected proteins that were greater than 2.0-fold enriched in Snx14-EGFP-APEX2 samples over negative controls lacking APEX2 (**Fig S3A, Circle 1 of Fig 3E**). Next, we focused on proteins annotated to localize to the ER network but also specific to the Snx14 interactome, since the ER was the primary organelle that manifested morphological alterations in *SNX14*^KO^ cells following palmitate exposure. We did this by eliminating proteins detected at similar levels in both the ER-APEX and Snx14-EGFP-APEX2 proteomics (**Fig S3A, B; Circle 2 of Fig 3E**). We then applied gene ontology enrichment analysis on the annotated ER-associated proteins specific to Snx14 (Circle 3 of Fig 3E, Fig S3C), revealing that ∼250 of these proteins are associated with cellular metabolism, and ∼100 with lipid metabolism (**Fig S3D**).

With this smaller candidate list, we compared them to a recently published genome-wide CRISPR/Cas9 screen identifying proteins whose loss sensitized cells to palmitate-induced lipotoxicity [46]. Remarkably, this unbiased screening identified Snx14 in the top 6% of proteins that were protective against palmitate (KO gene score: -2.28). This study also identified 310 proteins detected in our Snx14 interactome that exhibited a negative score greater than -1.8 for palmitate sensitivity [46] (**Supplementary Table 2; orange dots of Fig S3A; Circle 4 of Fig S3E**). These palmitate-sensitive proteins included FADS1, FADS2, SCD1, CEPT1 and HSD17B12 (**Fig S3F**).

As a final stage of analysis, we compared our list to our recently published interactome of Snx14 *Drosophila melanogaster* ortholog Snz, which also functions in lipid homeostasis in fruit fly adipocytes [26]. Snz proteomics identified the enzyme DESAT1, the major fly Δ-9 FA desaturase, as a key Snz functional interactor that was required for Snz-driven TG synthesis in *Drosophila* [26]. DESAT1 is the ortholog of human Δ-9 FA desaturase SCD1, which catalyzes the conversion of SFAs to MUFAs prior to their incorporation into glycerophospholipids or neutral lipids [47]. Indeed, several lines of evidence converged on SCD1 and its desaturation activity as being functionally linked to Snx14: 1) both proteins are ER-associated integral membrane proteins, and SCD1 was highly enriched in Snx14-EGFP-APEX2 proteomics in both *U2OS* and HEK293 cell lines (**Fig 3D, Fig S3A**); 2) SCD1 genetic or pharmacological perturbation hyper-sensitizes cells to palmitate-induced cell death, similar to Snx14 loss [13, 48, 49] (**Fig 3F**); 3) in genome-wide screening both Snx14 and SCD1 scored similarly in impact to palmitate sensitivity (SCD1: -2.42, Snx14: -2.28) [46]. Note that here, we are operationally defining a potential functional interaction between SCD1 and Snx14 based on their phonotypical similarities; this is distinct from any potential physical interaction. Based on this analysis, we chose to investigate the functional interplay between Snx14 and <u>SCD1-associated</u> FA desaturation in ER lipid homeostasis.

### *SNX14*^KO^ cell FA elevations are similar to cells with reduced SCD1 activity

To begin dissecting the functional interplay between Snx14 and FA metabolism, we conducted whole-cell lipidomic FA profiling. We pulsed WT and *SNX14*^KO^ cells for 2 hrs with 500μM palmitate, extracted polar lipids, and conducted gas chromatography (GC)-MS analysis to profile polar lipid fatty acyl chain length and saturation. Indeed, *SNX14*^KO^ cells exhibited a significant increase in total polar lipid-derived FAs (**Fig S4A**), as well as a significant increase in the abundance of 16-carbon length saturated fatty acyl chains (FA|16:0) in polar lipids relative to WT cells (**Fig 4A**). To determine whether this increase mimics alterations in polar lipids when SCD1 function is perturbed, we also treated WT cells with SCD1 inhibitor (SCD1i). Indeed, WT cells exposed to SCD1i exhibited similar increases in 16:0 saturated fatty acyl chains in their polar lipid fraction, mirroring elevations observed in *SNX14*^KO^ cells (**Fig 4A**). Furthermore, SCD1i-treated *SNX14*^KO^ cells exhibited no additional 16:0 acyl chain increases in the polar lipid profile, implying SCD1i treatment and Snx14 loss may perturb the same pathway.

**Figure 4:**
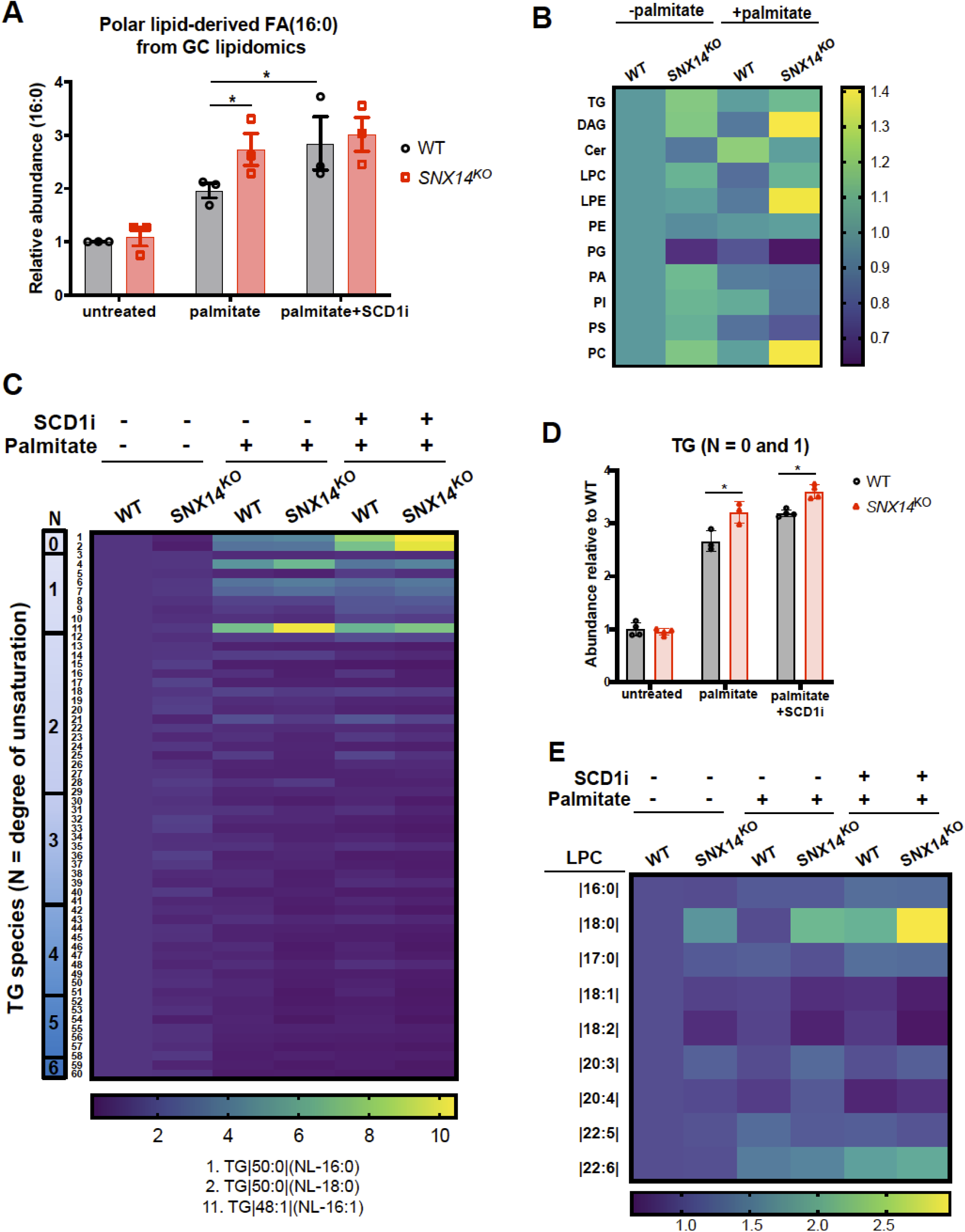
*SNX14*^KO^ cell lipidomic profiling is similar to cells with reduced SCD1 activity. **A**. Abundance of FAs (16:0) derived from polar lipids of WT and *SNX14*^KO^ cells relative to untreated WT under the following conditions – no treatment, palmitate treatment and treatment with palmitate and SCD1 inhibitor (SCD1i). Values represent mean ± SEM (n=3, *p<0.01, multiple t-test by Holm-Sidak method with alpha = 0.05). B. Heatmap indicating the relative change in abundance of individual lipid species of *U2OS* WT and *SNX14*^KO^ cells before and after palmitate treatment relative to untreated WT cells. C. Heatmap indicating the relative change in abundance of 60 different TG species of WT and *SNX14*^KO^ cells relative to untreated WT. These cells were either untreated or treated with palmitate or palmitate in presence of SCD1i. The label N indicates the number of double bonds in those group of TG species. The vertical serial number represents a different TG species where 1 is TG|50:0|(NL-16:0) and 11 is TG|48:1|(NL-16:1) and they exhibit the most changes in TG in the treated *SNX14*^KO^ cells. D. Abundance of TG (with 0 or 1 unsaturation) relative to untreated WT as analyzed in WT and *SNX14*^KO^ cells quantified from Fig 4C. Prior to lipid extraction and lipidomics, these cells were either untreated or treated with palmitate or palmitate in presence of SCD1i. Values represent mean±SEM (n=3, *p<0.01, multiple t-test by Holm-Sidak method with alpha = 0.05). E. Heatmap indicating the relative change in abundance of 9 different LPC species of WT and *SNX14*^KO^ cells relative to untreated WT. These cells were either untreated or treated with palmitate and palmitate in presence of SCD1i.

### TG, PC, and lysolipids lipids exhibit significant elevations in saturated acyl chains in *SNX14*^**KO**^ **cells**

To dissect how specific lipid classes are altered by Snx14 loss, we conducted global lipidomics via quantitative liquid-chromatography (LC)-MS analysis. WT and *SNX14*^KO^ cells were either left untreated or exposed to 2 hrs of palmitate, harvested, and analyzed. For comparative analysis, we again treated some cells with SCD1i. Importantly, this LC-MS analysis revealed fatty acyl properties such as chain length and saturation types of major lipid classes, including triglyceride (TG), diacylglyceride (DAG), phosphatidylethanolamine (PE), phosphophatidylcholine (PC), phosphatidic acid (PA), phosphatidylserine (PS), and lysophospholipids. Analysis revealed a significant (∼40%) increase in the levels of DAG, LPE and PC in *SNX14*^KO^ cells relative to WT cells following palmitate treatment (**Fig 4B**). However, relative abundances of most lipid classes between these two samples with or without palmitate treatment were not drastically altered.

Next, we examined the change in acyl chain saturation within each lipid class. In general, there was a greater proportion of saturated fatty acyl chains within several lipid classes. The most significant change in saturated fatty acyl chains in palmitate-treated *SNX14*^KO^ cells relative to WT was in TG and lyso-PC (LPC). Examining ∼60 different TG species revealed the most significant increase in TG species containing two or more saturated acyl chains. These TG species contained either 0 or 1 unsaturation among all three acyl chains (denoted as an N score of 0 or 1) (**Fig 4C**). For example, TG|48:1(NL-16:1), which contains one mono-unsaturated (16:1) acyl chain and two fully saturated acyl chains, was significantly elevated in *SNX14*^KO^ cells following palmitate treatment (**Fig 4C, species 11**). Notably, TG 50:0 (NL-16:0) and 50:0 (NL-18:0), both of which contained three saturated acyl chains, were also elevated in *SNX14*^KO^ cells compared to WT following palmitate addition and SCD1 inhibition, suggesting *SNX14*^KO^ cells incorporated SFAs into TG more than WT cells (**Fig 4C, species 1 and 2**). For more global analysis, we pooled the abundances of all TG species comprising only 0 or 1 total fatty acyl unsaturation. Indeed, TG pooling revealed that TG containing 0 or 1 unsaturation was significantly increased in palmitate treated *SNX14*^KO^ cells, and closely mirrored levels of WT cells treated with SCD1i (**Fig 4D**). Collectively, this suggests that the TG lipid profile of palmitate-treated *SNX14*^KO^ cells exhibits increased acyl chain saturation, and is similar to cells with decreased SCD1 function.

Among polar lipid species, the abundance of LPC species containing the saturated fatty acyl chain 18:0 was significantly increased in palmitate-treated *SNX14*^KO^ cells relative to WT (**Fig 4E**). In fact, levels of 18:0-containing LPC in *SNX14*^KO^ cells treated with palmitate closely mirrored WT cells treated with palmitate and SCD1i, again indicating that *SNX14*^KO^ cells closely resembled cells with inhibited SCD1 function by lipid profiling (**Fig 4E**). In line with this, there was a decrease in LPC species containing 18:1 or 18:2 acyl chains following palmitate treatment. LPE species containing 18:0 fatty acyl chains were also significantly increased in *SNX14*^KO^ cells even without palmitate treatment (**Fig S4B**). Profiling of PC, one of the most abundant glycerophospholipids, also revealed an increase in PC species with 18:0 fatty acyl chain in *SNX14*^KO^ cells, and that was more pronounced with palmitate treatment followed by SCD1 inhibition (**Fig S4D**). PS lipid profiles were similarly altered in *SNX14*^KO^, with increases in PS species containing 18:0 saturated fatty acyl groups, and a corresponding decrease in PS with 18:1 or 18:2 acyl chains (**Fig S4C**).

Collectively, LC-MS lipidomics suggests: 1) *SNX14*^KO^ cells exhibit elevated TG species containing two or more saturated acyl chains, 2) *SNX14*^KO^ cells have increased levels of LPC and LPE lysolipids containing saturated fatty acyl chains, 3) *SNX14*^KO^ cells exhibit more PC and PS with an 18:0 saturated acyl chain and less with 18:1 or 18:2, and 4) for TG and LPC, the lipid profiles of *SNX14*^KO^ cells exposed to palmitate closely mirror cells with inhibited SCD1 function.

### SCD1 activity can rescue *SNX14*^KO^ palmitate-induced lipotoxicity

Given the increased lipid saturation profiles of *SNX14*^KO^ cells, we next examined SCD1 protein levels in the context of Snx14 loss. SCD1 protein abundance was similar in WT and *SNX14*^KO^ cells under ambient conditions, but became elevated in both cell lines when exposed to 500μM palmitate. Notably, SCD1 protein levels were significantly more elevated in *SNX14*^KO^ cells exposed to palmitate. Snx14-Flag over-expressing cells returned SCD1 protein levels to WT levels, implying that *SNX14*^KO^ cells may further elevate SCD1 protein expression to compensate for Snx14 loss (**Fig 5A**,**B**).

**Figure 5:**
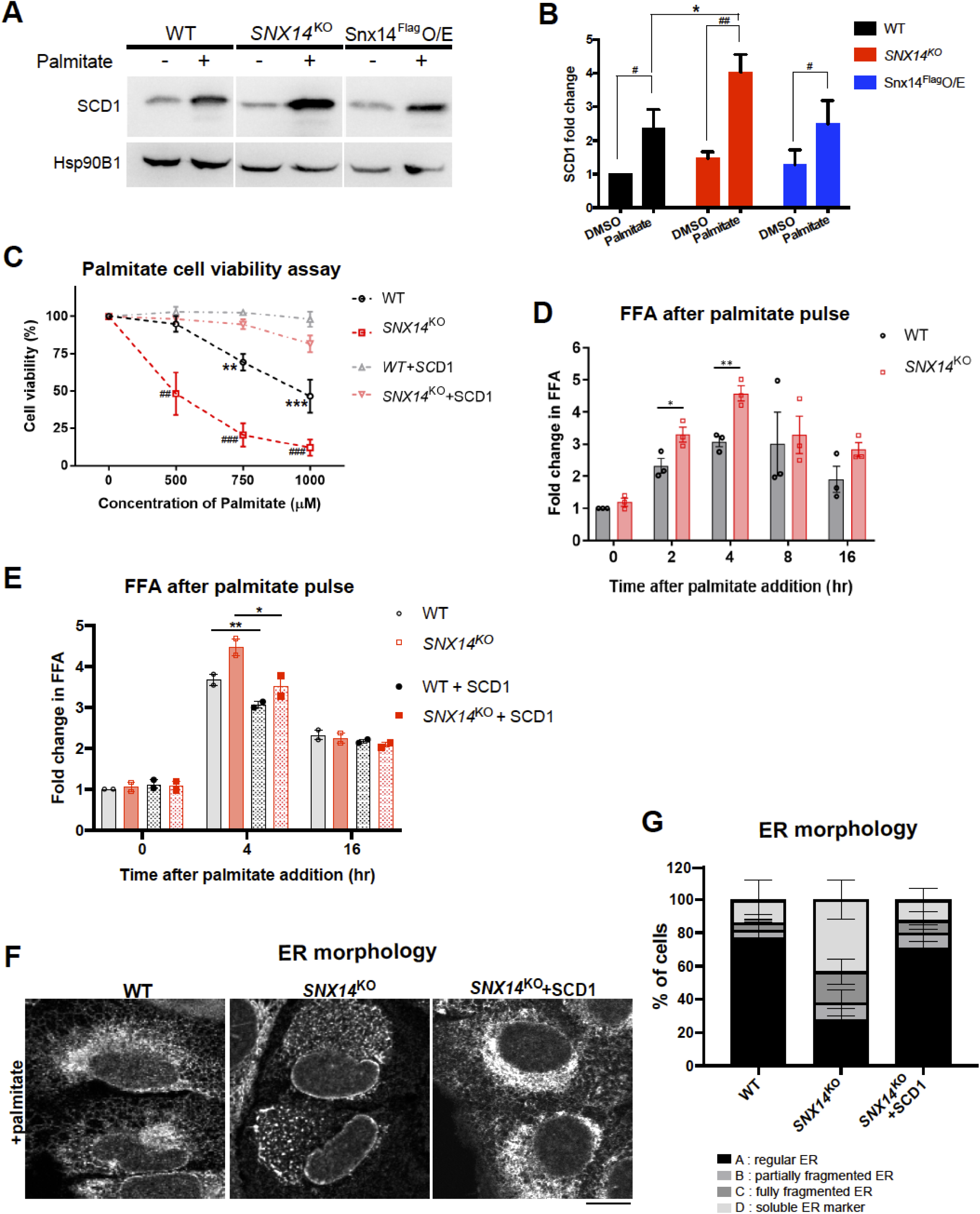
SCD1 activity can rescue *SNX14*^KO^ palmitate-induced lipotoxicity. **A**. Western blot of SCD1 and Hsp90B1 (ER marker) before and after overnight palmitate treatment in WT, *SNX14*^KO^ and Snx14Flag over-expressed (O/E) cells. **B**. Ratio of the intensity of the protein bands of SCD1 over Hsp90B1 are quantified from Fig 5A, and plotted as fold change relative to untreated WT whose ratio is set as 1. Values represent mean ± SEM. Significance test between before and after palmitate treatment denoted as # (n=3, ^#^p<0.1, ^##^p<0.001, multiple t-test by Holm-Sidak method with alpha = 0.05). Significance test between WT, *SNX14*^KO^ and Snx14FlagO/E is denoted as *(n=3, *p<0.01, multiple t-test by Holm-Sidak method with alpha = 0.05) **C**. Cell viability (%) of WT and *SNX14*^KO^ cells, showing sensitivity of both WT and *SNX14*^KO^ cells are rescued with overexpression of SCD1 following addition of increasing concentration (0, 500, 750, 1000μM) of palmitate for 2 days. The assay was repeated thrice in triplicates. Values represent mean±SEM. Significance test between WT and WT+SCD1 denoted as * (n=3, **p<0.001, ***p<0.0001, multiple t-test by Holm-Sidak method with alpha = 0.05) Significance test between *SNX14*^KO^ and *SNX14*^KO^+SCD1 denoted as ^#^ (n=3, ^##^p<0.001, ^##^ p<0.0001,multiple t-test by Holm-Sidak method with alpha = 0.05). **D**. TLC of neutral lipids and FFAs performed in WT and *SNX14*^KO^ cells treated with 500μM palmitate for 0, 2, 4, 8, 16 hours. Quantification of relative fold change in FFA (normalized to cell pellet weight) with respect to untreated WT from this TLC. Values represent mean±SEM (n=3, *p<0.01, multiple t-test by Holm-Sidak method with alpha = 0.05). **E**. TLC of neutral lipids and FFAs performed in WT and *SNX14*^KO^ cells before and after overexpression of SCD1 following exposure to 500μM palmitate for 0, 4, 16 hours. Quantification of relative fold change in FFA (normalized to cell pellet weight) with respect to untreated WT from this TLC. Values represent mean±SEM (n=3, *p<0.01, **p<0.001, multiple t-test by Holm-Sidak method with alpha = 0.05). **F**. Immunofluorescent (IF) labeling of the ER with α-HSP90B1 (ER marker) antibody in *SNX14*^KO^ cells before and after overexpression of SCD1 following overnight palmitate treatment. Scale bar = 10μm. **G**. Percentage of palmitate treated WT and *SNX14*^KO^ cells quantified and grouped based on whether the ER morphology is regular (A), partially fragmented (B), fully fragmented (C) or completely soluble (D). Total ∼100 cells quantified from 3 experiments. Values represent mean±SEM.

Since *SNX14*^KO^ cells exhibited slightly elevated SCD1 levels, we queried whether ectopic SCD1 over-expression could reduce *SNX14*^KO^ associated palmitate hyper-sensitivity. Strikingly, SCD1 over-expression rescued *SNX14*^KO^ cell viability, and *SNX14*^KO^ cells now responded similarly to WT cells exposed to dose-dependent palmitate treatment (**Fig 5C**). *SNX14*^KO^ cells were also rescued by exposure to mixtures of palmitate and oleate, the MUFA and product of SCD1 enzymatic activity (**Fig S5A**). Since *SNX14*^KO^ cells also displayed defective LD morphology with palmitate, we tested whether SCD1 over-expression could restore LD levels. Indeed *SNX14*^KO^ cells over-expressing SCD1 manifested more LDs following palmitate exposure (Fig S2A,B). Since LDs are lipid reservoirs, we also determined whether this SCD1-mediated rescue required the incorporation of FAs into TG for LD storage. We exposed *SNX14*^KO^ cells to DGAT1/2 inhibitors (DGATi) in the presence of palmitate, and monitored cell viability. Surprisingly, SCD1 over-expression rescued *SNX14*^KO^ cells even with DGATi, suggesting this SCD1-mediated rescue functions upstream of TG synthesis (**Fig S5B**).

Since *SNX14*^KO^ and SCD1i-exposed cells displayed some lipidomics similarities, and SCD1 over-expression mitigated palmitate-induced *SNX14*^KO^ cell death, we hypothesized that *SNX14*^KO^ cells had defects processing SFAs. To test this, we exposed WT and *SNX14*^KO^ cells to 500μM palmitate for 0, 2, 4, 8, and 16 hr, extracted whole cell lipids and conducted thin layer chromatography (TLC) to monitor changes in free FAs (FFAs) and neutral lipids. As expected, both WT and *SNX14*^KO^ cells exhibited elevated FFA levels following 2 and 4 hrs palmitate exposure, but *SNX14*^KO^ exhibited significantly elevated FFAs compared to WT (**Fig 5D, S5C**). SCD1 over-expression reversed this FFA elevation at 4 hrs in *SNX14*^KO^ cells, implying the elevated FFA pool was composed of SFAs that could be processed by SCD1 (**Fig 5E, S5D**). As a key control, we monitored uptake of radio-labeled ^14^C-palmitate in WT and *SNX14*^KO^ cells, to test whether elevated FFA accumulation in *SNX14*^KO^ cells was due to increased FA uptake. We confirmed that FA uptake was not altered by Snx14 loss (**Fig S5E**). This indicates that *SNX14*^KO^ cells accumulate FFAs following palmitate uptake, consistent with a defect in palmitate processing, and these effects can be reversed by SCD1 over-expression.

To understand whether the ER fragmentation observed in *SNX14*^KO^ cells is associated with defects in palmitate processing, we examined ER morphology in SCD1 over-expressed *SNX14*^KO^ cells following palmitate exposure. Remarkably, ectopic expression of SCD1 in palmitate-treated *SNX14*^KO^ cells rescued ER morphology (**Fig 5F, G**). Collectively, this suggests that SCD1 over-expression can rescue *SNX14*^KO^ elevated lipid saturation, LD morphology, and ER fragmentation.

### Snx14 functionally interacts with SCD1 in the ER network

Since Snx14 and SCD1 appeared to have phonotypical similarities, and SCD1 over-expression could rescue aspects of Snx14 loss, we next examined whether the two proteins could co-immunoprecipitate (co-IP), which would suggest their close proximity to each other within the ER network. We generated *U2OS* cell lines stably expressing either 3xFlag-tagged Snx14 or GFP, conducted anti-Flag immuno-precipitation, and Western blotted the IP lysates for endogenous SCD1. SCD1 was detected in the Snx14-Flag sample but not GFP-Flag (**Fig 6A**). Intriguingly, Snx14-Flag co-IPed endogenous SCD1 both with and without palmitate addition (**Fig 6A**). To determine whether this interaction was specific, we Western blotted for PLIN3 which was highly enriched in the Snx14 APEX2-proteomics (**Fig 3D, Fig S3A**). Notably, PLIN3 was not detected in the Snx14 co-IP lysate (**Fig S6A**).

**Figure 6:**
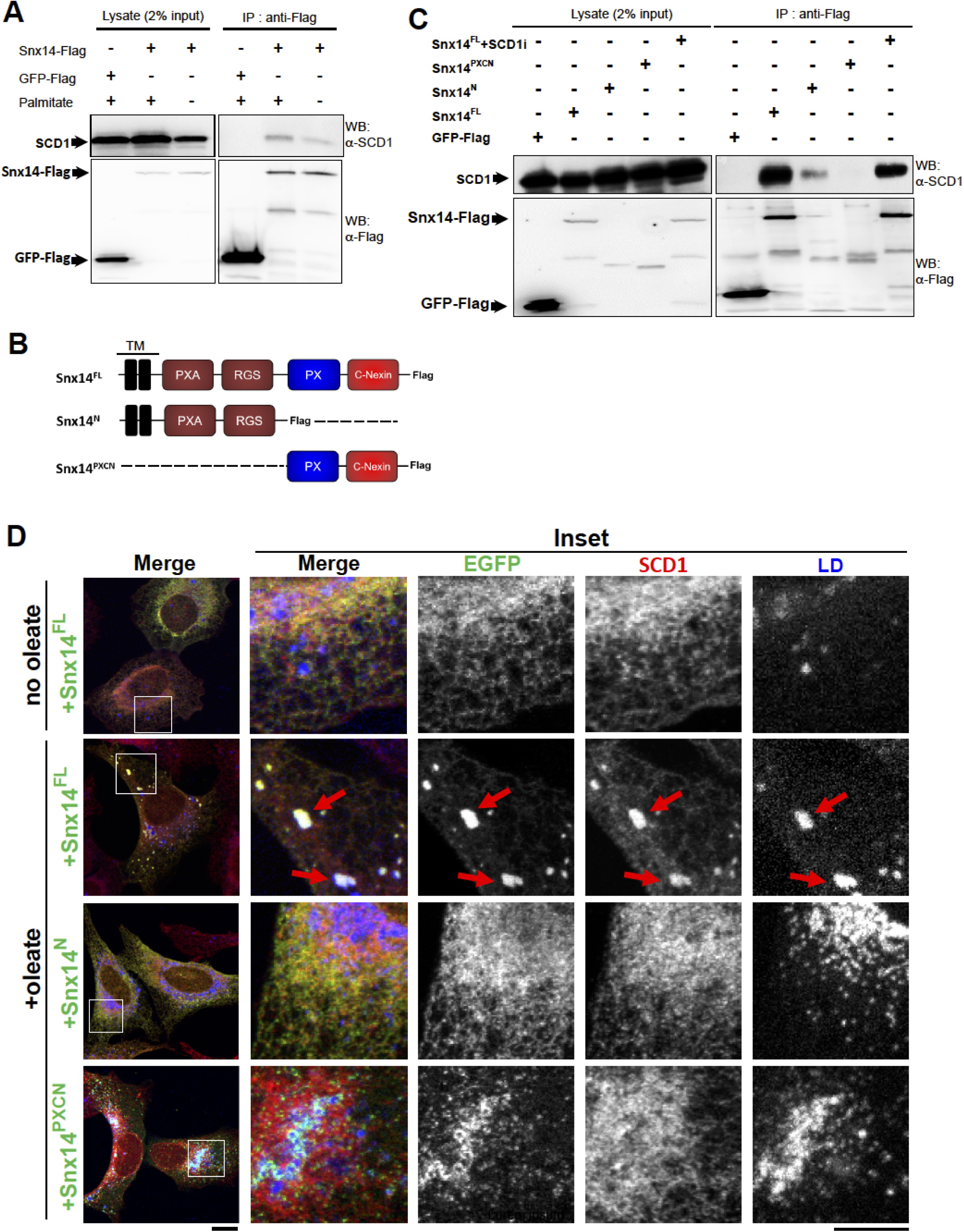
Snx14 interacts with SCD1 in the ER network. A. Western blotting with anti-Flag and anti-SCD1 antibody of 2% input lysate from GFP-Flag and Snx14-Flag expressing cells with and without palmitate treatment reveals relative expression of GFP-Flag, Snx14-Flag and SCD1. Co-immunoprecipitation (Co-IP) of SCD1 with Flag tagged constructs reveals presence of SCD1 in Snx14-Flag and not GFP-Flag enriched beads when western blotted with anti-Flag and anti-SCD1 antibody. B. Schematic diagram of Snx14 fragments C-terminally tagged with either 3X Flag or EGFP. Snx14^FL^ depicts the full length human Snx14. Snx14^N^ is the N-terminal fragment that spans from the beginning and includes TM, PXA and RGS domains. Snx14^PXCN^ is the C-terminal half including the PX domain and C-Nexin domains. **C**. Lanes represent 2% input and IP from GFP-Flag, all 3XFlag tagged Snx14 constructs (Snx14^FL^, Snx14^N^, Snx14^PXCN^) and SCD1i treated Snx14^FL^-3XFlag expressing U2OS cells. Western blotting with anti-Flag and anti-SCD1 antibody reveals relative expression of all the Flag tagged constructs and SCD1 in all these samples. **D**. IF staining of U2OS cells expressing Snx14^FL^, Snx14^N^, Snx14^PXCN^ with anti-EGFP(green), anti-SCD1(red) antibody and imaged with confocal microscope. LDs were stained with MDH (blue). The cells were either untreated or treated with oleate. Scale bar = 10 μm.

Next we dissected what regions of Snx14 were sufficient to co-IP SCD1. We used cell lines stably expressing the Flag-tagged N-terminal half of Snx14 encoding the transmembrane (TM), PXA, and RGS domains (Snx14^N^), or a C-terminal half with the PX and C-Nexin domains (Snx14^PXCN^) (**Fig 6B**). Co-IP experiments revealed that Snx14^N^, but not Snx14^PXCN^, was sufficient to pull down SCD1 (**Fig 6C**). To test whether this Snx14:SCD1 co-IP required SCD1 desaturase activity, we conducted the co-IP with full length Snx14 (Snx14^FL^) in the presence of SCD1i [50]. Inhibited SCD1 still co-IPed with Snx14^FL^, indicating this interaction did not require SCD1 enzymatic activity (**Fig 6C**).

Given Snx14 and SCD1 could co-IP *in vitro*, we tested whether this interaction occurred in intact cells. We exploited the previous observation that Snx14 enriches at ER-LD contacts following oleate addition [23]. Given this, we queried whether Snx14 over-expression was sufficient to drive the accumulation of SCD1 at ER-LD interfaces, since it normally resides throughout the ER network. As expected, immunofluorescence (IF) labeling of endogenous SCD1 in non-oleate treated cells transfected with Snx14-EGFP revealed SCD1 co-localization with Snx14 throughout the ER network (**Fig 6D, S6B**). Following oleate addition, Snx14-EGFP accumulated at ER-LD contacts, and co-localized with SCD1 foci which also enriched at these sites (**Fig 6D, red arrows, S6B**). In contrast, expression of Snx14^PXCN^-EGFP, which accumulates around LDs [23], failed to colocalize with SCD1 foci, consistent with Snx14^PXCN^ being insufficient to co-IP SCD1 (**Fig 6D, S6B**). Snx14^N^-EGFP localized throughout the ER network following oleate, since it lacks the previously identified LD-targeting region in the CN domain [23], and this co-localized with SCD1 throughout the ER network but did not promote SCD1 foci at ER-LD contacts, consistent with co-IP results (**Fig 6D, S6B**). Collectively, this suggests that Snx14 co-IPs SCD1 from cell lysates, and can interact with SCD1 in cells. It should be noted that the ER-LD SCD1 accumulations we observe require Snx14 over-expression; we do not detect SCD1 enrichment at ER-LD contacts under more ambient conditions.

### Snx14 loss does not impact SCD1 enzymatic activity, but Snx14 requires its FA-binding PXA domain for function

To further dissect how Snx14 and SCD1 contribute to SFA metabolism at the ER, we examined whether Snx14 regulates SCD1 enzymatic activity. We directly assayed SCD1 activity *in vitro* via a well-established radio-label based desaturase assay [10]. We pulsed microsomal fractions isolated from WT or *SNX14*^KO^ cells with (9,10-^3^H)-stearoyl-CoA, a SCD1 substrate. SCD1 activity releases free ^3^H when stearoyl-CoA is mono-unsaturated, which can be directly detected by scintillation counting. As a positive control, we treated samples with SCD1i, and detected a significant decrease in free ^3^H indicating reduced SCD1 activity (**Fig 7A**). However, there was no significant change in relative SCD1 activity in *SNX14*^KO^ cells, suggesting Snx14 is not required for SCD1 activity.

**Figure 7:**
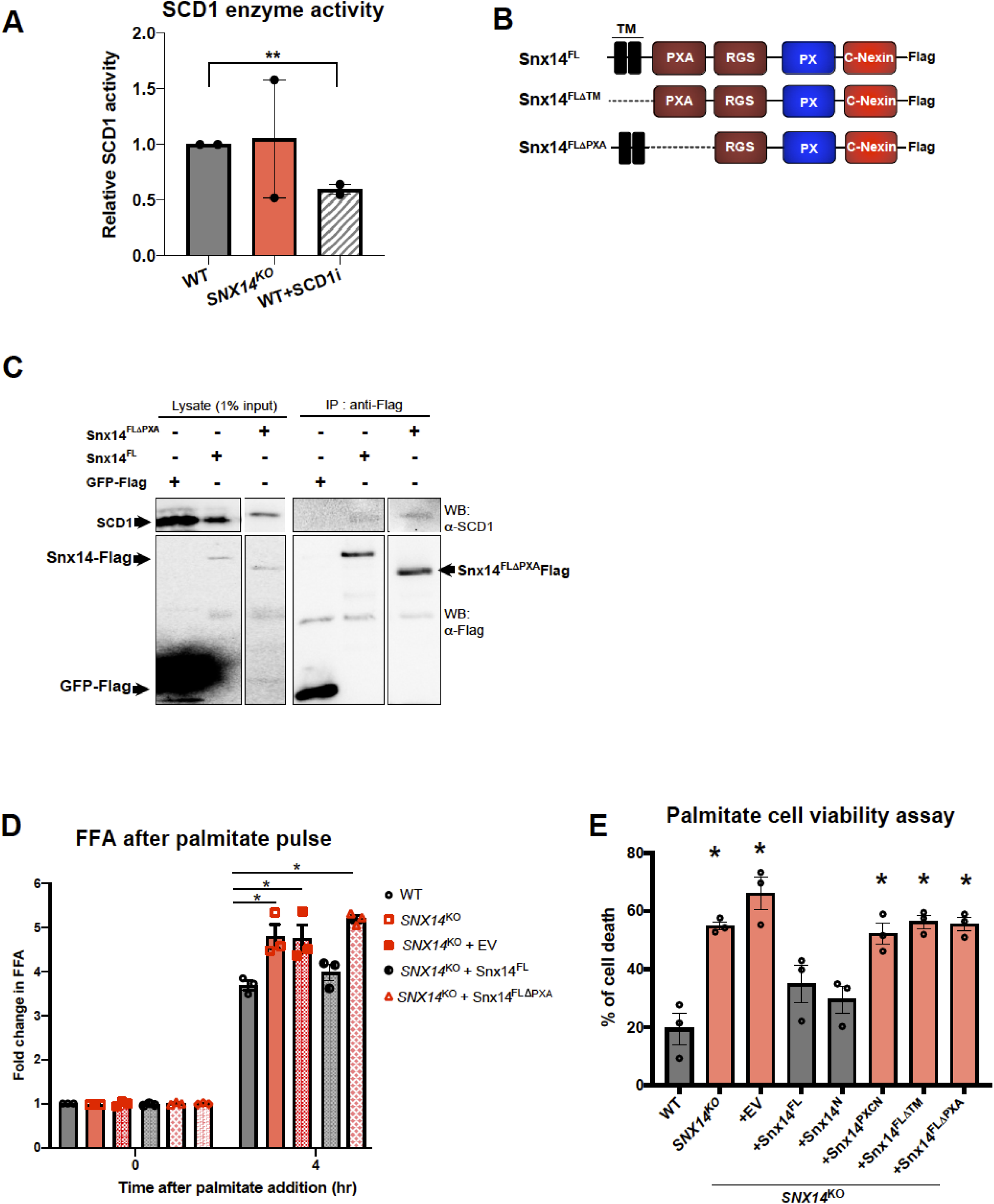
Snx14 loss does not impact SCD1 enzymatic activity, but Snx14 requires FA-binding PXA domain for function. A. SCD1 enzyme activity quantified in WT, SNX14-KO and SCD1i-treated WT cells relative to WT cells. Values represent mean±SEM (n=2, **p<0.001, multiple t-test by Holm-Sidak method with alpha = 0.05). B. Schematic diagram of Snx14 fragments C-terminally tagged with 3X Flag. Snx14^FL^ depicts the full length human Snx14. Snx14^FLΔPXA^ and Snx14^FLΔTM^ are the full length Snx14 excluding the PXA domain and TM domain respectively. C. Lanes represent 1% input and IP from GFP-Flag, 3XFlag tagged Snx14 constructs (Snx14^FL^, Snx14^FLΔPXA^, Snx14^FLΔTM^) expressing U2OS cells. Western blotting with anti-Flag and anti-SCD1 antibody reveals relative expression of all the Flag tagged constructs and SCD1 in all these samples. Snx14^FLΔPXA^ could co-IP SCD1 similar to Snx14^FL^ whereas Snx14^FLΔTM^ could not pull down SCD1. D. Quantification of fold change in FFA (normalized to cell pellet weight) relative to untreated WT from TLC of whole cell neutral lipids extracted from WT, *SNX14*^KO^, *SNX14*^KO^ cells expressing either EV, Snx14^FL^, or Snx14^FLΔPXA^ which are either untreated or treated with palmitate for 4hr. Values represent mean±SEM (n=3, *p<0.01, multiple t-test by Holm-Sidak method with alpha = 0.05). E. Cell death (%) after exposure to 500μM of palmitate in WT, *SNX14*^KO^, and on re-addition of empty vector (EV), Snx14^FL^, Snx14^N^, Snx14^PXCN^, Snx14^FLΔTM^, Snx14^FLΔPXA^ to *SNX14*^KO^. Values represent mean±SEM. Significance test compared with WT is denoted as * (n=3, *p<0.01, multiple t-test analysis by Holm-Sidak method with alpha = 0.05.

Since Snx14 yeast ortholog Mdm1 had previously been shown to directly bind to FAs *in vitro* via its PXA domain [25], we next interrogated whether Snx14 requires its PXA domain for function. We generated cell lines stably expressing Snx14 lacking its PXA domain (Snx14^FL^ΔPXA) (**Fig 7B)**, and queried whether this construct could co-IP with SCD1 as well as process the accumulated FFAs in *SNX14*^KO^ cells following palmitate exposure. Intriguingly, Snx14^FL^ΔPXA could co-IP with SCD1 (**Fig 7C**). However, in contrast to *SNX14*^KO^ cells expressing full length Snx14 (Snx14^FL^) which could had normal FFA levels, *SNX14*^KO^ expressing Snx14^FL^ΔPXA cells exhibited elevated FFA levels similar to *SNX14*^KO^ cells. This indicates that the Snx14 PXA domain was required for FFA processing (**Fig 7D**). Importantly, here also we confirmed that FA uptake was not altered by expressing Snx14^FL^ or Snx14^FL^ΔPXA in *SNX14*^KO^ cells, indicating that the elevated FFA accumulation in *SNX14*^KO^ cells expressing Snx14^FL^ΔPXA was not due to increased FA uptake (**Fig S5E**).

Next, we assayed whether the PXA domain was necessary to rescue *SNX14*^KO^ cell viability following palmitate exposure. Indeed, Snx14^FL^ΔPXA failed to rescue *SNX14*^KO^ cell viability, indicating the PXA was required for Snx14-mediated protection from palmitate (**Fig 7E**). However, *SNX14*^KO^ cells expressing Snx14^N^ were protected from palmitate, suggesting this minimal Snx14 fragment which contains an ER-anchored PXA domain is sufficient for palmitate protection. However, a Snx14 construct which lacked the N-terminal TM domain but encoded all other domains (Snx14^FL^ΔTM) could not rescue the *SNX14*^KO^ cell viability, indicating that TM-mediated ER association is also required for Snx14 function (**Fig 7B, E**). Collectively, this suggests that Snx14 loss does not impact SCD1 enzymatic activity *in vitro*, but implies that the ER-associated Snx14 PXA domain may interact with FAs at or near the ER network in a manner that promotes SCD1-mediated FA processing.

## Discussion

FAs are essential cellular components that act as energy substrates, membrane components, and key signaling molecules. These functions are dependent on the chemical features of FAs such as their chain length and saturation degree, which influence membrane fluidity and impact organelle structure and function [10, 46]. Lipid homeostasis depends on FA processing and channeling to specific organelle destinations. When cells experience elevated intracellular FA levels, they respond by increasing FA processing and storage. Excess FAs are incorporated into neutral lipids and stored in LDs, which protect cells from lipotoxicity [51]. Much of the machinery to achieve this resides at the ER, and proper ER-LD crosstalk by proteins such as seipin and the FATP1-DGAT2 complex is essential to maintain lipid homeostasis [17-19]. Our earlier work revealed a role for Snx14 in ER-LD crosstalk, the loss of which contributes to the cerebellar ataxia disease SCAR20 for unknown reasons [23]. Here, we report the proteomic composition of Snx14-associated ER-LD contacts, and provide mechanistic insights into the function of Snx14 in FA metabolism. We find that Snx14 loss impacts the ability of cells to maintain proper lipid saturation profiles. In line with this, following SFA exposure *SNX14*^KO^ cells display defects in ER morphology, FFA accumulation and elevated SFA incorporation into membrane glycerophospholipids, lyso-phospholipids, and TG, which ultimately impacts cell homeostasis.

Utilizing APEX2-based proteomics analysis, we reveal the protein composition of Snx14-positive ER-LD contacts formed during oleate treatment. Notably, these contacts contain proteins associated with the LD surface such as PLIN2, PLIN3, and PNPLAP2. We also detect numerous proteins involved in FA processing (ACSL4, SCD1) and lipid/sterol biosynthesis (LPCAT1, LPCAT3, SOAT1), indicating that ER-LD contacts act as lipogenic ER sub-domains for several lipid species, as well as hotspots for FA processing. These proteomics reveal many proteins previously identified through a similar APEX-based proteomics study which targeted APEX to the LD surface, providing consistency in the method, as well as underscoring the tight functional association between LDs and the ER network [44]. Among these proteins, we focused on investigating the functional interplay between Snx14 and SCD1-associated SFA metabolism, which provided important insights in understanding the hypersensitivity of *SNX14*^KO^ cells to palmitate exposure. Although we cannot conclude that Snx14 and SCD1 directly interact, their phenotypical similarities and functional interplay provide important insights that may help in the therapeutic treatment of SCAR20.

Through lipidomic analysis and biochemistry, we show that Snx14 loss phenotypically mimics aspects of the enzymatic inhibition of Δ-9-FA desaturase SCD1, which catalyzes the oxidization of SFAs to MUFAs (**Fig 8**) [47]. In line with this, SCD1 over-expression can rescue palmitate sensitivity in Snx14-deficient cells. We also find that SCD1 and Snx14 can co-IP from cell lysates, and that Snx14 over-expression can promote SCD1 enrichment at ER-LD contacts under distinct metabolic conditions. The data presented are consistent with a model where Snx14 may promote FA processing to maintain ER homeostasis. One possibility is that Snx14 acts as an organizational scaffold for enzymes including SCD1 within the ER network, enabling FAs to be processed. In agreement with this, Snx14 requires its FA-binding PXA domain to function, implying that Snx14 could bind FA substrates and present them to SCD1 during periods of elevated FA influx. However, it should be noted that *SNX14*^KO^ cells do not display altered SCD1 enzymatic activity, indicating that Snx14 is not a direct enzymatic regulator of SCD1 *per se*, nor required for its activity. We also cannot rule out the possibility that Snx14 may interact or present FAs to other enzymes in the ER network, which could explain other alterations in the lipidomics profile of *SNX14*^KO^ cells.

**Figure 8:**
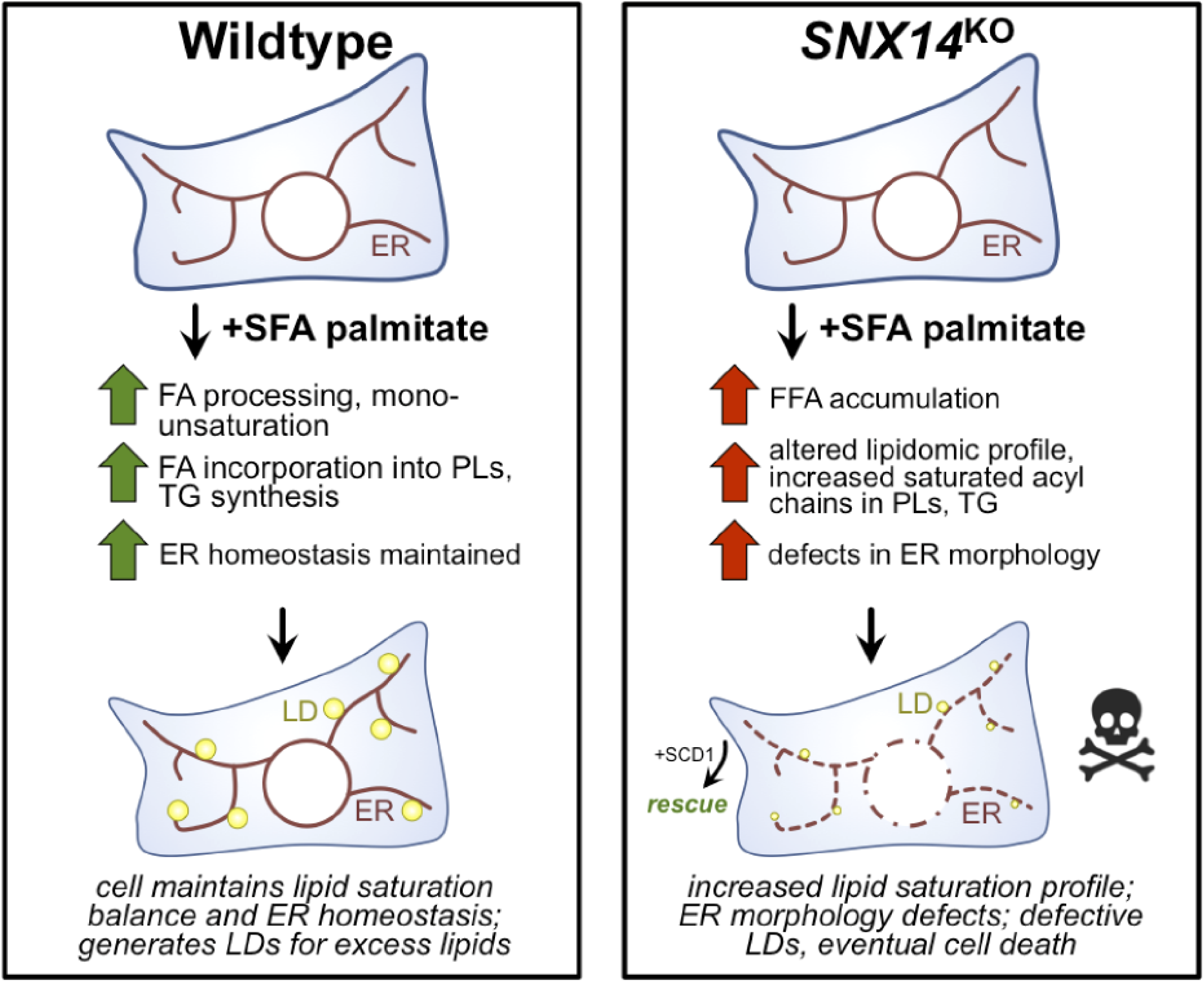
Working model for Snx14 in maintaining ER lipid homeostasis when treated with excessive SFAs.

Snx14 is highly conserved in evolution, and related studies of Snx14 orthologs provide mechanistic insights into Snx14’s role in lipid homeostasis. *Drosophila* ortholog Snz functionally interacts with SCD1 ortholog DESAT1 in the adipocyte cell periphery, a sub-cellular region rich in FA processing [26]. Similarly, yeast ortholog Mdm1 functions in FA activation and LD biogenesis near the yeast vacuole [25, 52]. As such, Mdm1-deficient yeast are sensitive to high dosages of exogenous FAs. Studies of both Snz and Mdm1 also reveal these proteins localize to specific sub-regions of the ER network, and thereby help to demarcate ER sub-domains through their inter-organelle tethering capabilities. Thus, an emerging possibility is that Snx14 family proteins act as organizational scaffolds within the ER network, and recruit enzymes into ER sub-domains to more efficiently process lipids.

Collectively, our observations provide a framework for understanding how Snx14 loss contributes to the cerebellar ataxia disease SCAR20. A growing number of neurological diseases are attributed to loss of proteins that function in ER-localized lipid metabolism or in the maintenance of ER architecture [27-29]. Our data are consistent with a model where Snx14 loss perturbs the ability of cells to maintain FA metabolism and membrane lipid composition, ultimately elevating SFA levels and saturated fatty acyl chain incorporation in membrane lipids. Such alterations will ultimately effect membrane fluidity, organelle function, and contribute to cellular lipotoxicity and the progressive death of cells like neurons, a pervasive symptom of SCAR20 disease [20, 22]. Additionally, although we focused our analysis here on the interplay between Snx14 and SFA metabolism, the APEX2-based proteomics and lipidomics analysis will provide insights into understanding proteins and lipogenic reactions at ER-LD contacts.

## Materials and Methods

### Cell culture

U2OS and HEK293 cells were maintained and expanded in DMEM (Corning, 10-013-CV) media containing 10% Cosmic Calf Serum (Hyclone, SH30087.04) and 1% Penicillin Streptomycin Solution (100X, Corning, 30-002-Cl). All cells were cultured at 37°C and 5% CO2. The cells were passaged with Trypsin-EDTA (Corning, 25-053-Cl) when they reach 80-90% confluency. For the cell viability assay, cells were treated with different concentration of FFA (250, 500, 750, 1000 μM) for 48 hours. For all other FFA treatment, the cells were incubated with 500μM of either oleate or palmitate for the indicated period of time. In all the experiments the FFAs were conjugated with fatty acid free BSA (Sigma, A3803) in the ratio of 6:1.

### Chemicals and reagents

For cell treatments, the following reagents were used for indicated periods of time – (1) SCD inhibitor (Abcam, ab142089) – 4 μM in DMSO (2) Myriocin (Sigma, M1177) – 50 μM in DMSO (3) Tunicamycin (Sigma) – 5 μg/ml in DMSO for 6hr (4) Etomoxir (Cayman chemical, 11969) – 10 μM in DMSO (5) DGAT inhibitors include DGAT1 inhibitor (A-922500, Cayman chemical, 10012708) – 10 μM in DMSO, and DGAT2 inhibitor (PF-06424439, Sigma, PZ0233)– 10 μM in H_2_O (6) IRE1 inhibitor 4μ8c (Sigma, SML0949) – 64μM in DMSO (7) PERK inhibitor I, GSK2606414 (EMD Millipore, 516535) - 30 nM in DMSO (c) Caspase 6/8 inhibitor (Sigma, SCP0094) - 40μM in DMSO.

### Generation of stable cell line using lentiviral transduction

Stable cell line – U2OS cells stably expressing all C-terminally 3X Flag tagged SNX14^FL^, SNX14^N^ and SNX14^PXCN^ generated in the previous study (ref) has been used here. Both Snx14^FL^ΔTM and Snx14^FL^ΔPXA, with 3XFlag tag at the C-terminal, were generated following PCR amplification using Snx14^FL^-3XFlag (primers available on request) and cloning into the plvx lentiviral vector with puromycin (puro) resistance cassette. Similarly GFP-Flag plasmid was cloned into plvx-puro vector. The cloned plasmids and lentiviral packaging plasmids were transfected together into 293T cells to generate lentivirus which were transduced into U2OS cells. Puromycin was used to select and expand the transduced cells expressing the plasmid. The generated stable cell lines were stored in liquid nitrogen before their use in experiments.

Stable U2OS cells expressing Snx14-EGFP-APEX2 was used from [23]. Similar stable cell line generation method as described above was used to generate EGFP-APEX2-Sec61β which is also puromycin resistant. cDNA library was used for PCR amplification of Sec61β. *SNX14*^KO^ *U2OS* cells used here was generated using CRISPR/Cas9 in [23].

### Cloning and transient transfection

All C-terminally EGFP tagged Snx14 constructs i.e. SNX14^FL^, SNX14^N^ and SNX14^PXCN^ used were generated in [23]. Snx14-EGFP-APEX2 was cloned in pCMV vector for transient transfection. The primers used for cloning are available on request. HEK293 and U2OS cells were transiently transfected using PEI-Max Transfection reagent (Polysciences, 24765) and optimem (gibco, 31985-070) for 48 hours prior to experiments.

### Cell viability assay

The cell viability assay protocol was adapted from [30, 31]. Cells were seeded at ∼40% confluency and maintained in cell culture media overnight in a 12 well plate. On the following day, the cells were treated with indicated concentration of FFA for 2 days. Then the cells were washed with PBS, fixed with 4% PFA, stained with 0.1% crystal violet (Sigma) for 30 mins, excess stain was washed, and then the stain from the surviving cells were extracted with 10% acetic acid, whose optical density (OD) was measured at 600nm. The percentage of survival cells (cell viability) were quantified relative to the untreated cells whose OD measurement is set at 100% survival. The assay was performed thrice in triplicates.

### Electron Microscopy

The U2OS cells were cultured under desired condition on MatTek dishes and processed in the UT Southwestern Electron Microscopy Core Facility. The cells were fixed with 2.5% (v/v) glutaraldehyde in 0.1M sodium cacodylate buffer, rinsed thrice in the buffer, and post-fixed in 1% osmium tetroxide and 0.8 % K_3_[Fe(CN_6_)] in 0.1 M sodium cacodylate buffer at room temperature for 1 hr. After rinsing the fixed cells with water, they were en bloc stained with 2% aqueous uranyl acetate overnight. On the following day, the stained cells were rinsed in buffer and dehydrated with increasing concentration of ethanol. Next they were infiltrated with Embed-812 resin and polymerized overnight in a 60°C oven. A diamond knife (Diatome) was used to section the blocks on a Leica Ultracut UCT (7) ultramicrotome (Leica Microsystems), which were collected onto copper grids, and post stained with 2% aqueous uranyl acetate and lead citrate. A Tecnai G^2^ spirit transmission electron microscope (FEI) equipped with a LaB_6_ source and a voltage of 120 kV was used to acquire the images.

### Immunofluorescent (IF) staining

For IF staining the cells were fixed with 4% PFA in PBS for 15-30 mins at room temperature. Then the cells were washed with PBS and permeabilized with 0.2% NP40 in PBS at RT for 3 mins. The cells were incubated with IF buffer (PBS containing 3% BSA, 0.1% NP40 and 0.02% sodium azide) for 40 mins to block non-specific binding. Next the cells were stained with primary antibody in IF buffer for 1 hr, washed with PBS thrice and stained with secondary antibody in IF buffer for 30 mins, again washed thrice with PBS, and used for imaging.

The primary antibodies used in this study are mouse anti-Hsp90B1 (1:300; Sigma, AMAb91019), rabbit anti-EGFP (1:300; Abcam, ab290), mouse anti-EGFP (1:200, Abcam, ab184601), rabbit anti-SCD1 (1:100; Abcam, ab39969), streptavidin-alexa 647 (Thermofisher Scientific, S21374). The secondary antibodies are donkey anti-mouse AF488 (Thermo Fisher, A21202) and donkey anti-rabbit AF594 (Thermo Fisher, A21207) used at a dilution of 1: 1000. LDs were stained with MDH (1:1000; Abgent, SM1000a) for 15 mins.

### Confocal microscopy and image analysis

A Zeiss LSM 780 Confocal Microscope was used to acquire the images of the cells with a 63X oil immersion objective. Approximately 4-5 Z-sections of each image were taken. ImageJ was used to analyse the images and represent them. The images were merged by max z-projection prior to analysis.

Analysis of ER morphology – We quantified the ER morphology from 3 sets of experiments, imaging approx. 6 cells in one field of view and six fields of view in each experiment. This gave us ∼100 cells to quantify the ER morphology which is grouped into four classes – (A) regular and intact ER (B) ER with partial fragmentation where the nucleus is still intact (C) fully fragmented ER, with distinct bulges where the nuclear envelope also disappeared, (D) the ER stain is soluble. Examples of A,B,C,D is shown in Fig 2. For each set of experiment, percentage of cells of each class of ER morphology is quantified and plotted.

Quantification of LDs – Using ImageJ, the images with MDH stained LDs were grayscaled and inverted (Fig S2A). The cell boundary was traced by a freehand line tool. To analyse the average area covered by LDs per cell, the following steps in ImageJ were used – 1) LDs were processed by using ‘Subtract background’ in ‘Process’. Next in ‘Process’, we used ‘Filter’ to apply ‘Gaussian blur’. To get rid of the background, we then subtracted the blurred image from the original in ‘image calculator’ in ‘Process’. 2) We ‘threshold’-ed the modified image and then applied ‘analyse particles’. The LDs were clustered in these images and could not be distinguished from one another. So, we analysed the average area covered by LDs per cell, which provided an estimate of both size and number of LDs.

### RNA extraction and qRT-PCR

To extract RNA, tissue culture cells were solubilized in TRIzol and collected from dishes, then treated with chloroform and centrifuged at 12000g for 15mins. To precipitate RNA, the upper colorless aqueous layer was collected, and isopropanol was added. Next, the supernatant was discarded and the RNA precipitate was washed by ethanol, air-dried and resuspended in RNase-free water. This RNA was used to generate cDNA using Bio-Rad commercial kit (iScript cDNA synthesis kit 1708891). The cDNA was used for qPCR reaction to analyse spliced xbp1 (s-xbp1) mRNA level which is an indicator of UPR activity that results from ER stress. Cells treated with tunicamycin (5μg/ml) for 6 hr was used as a control for UPR activity. The internal control used for the qPCR reaction was β-actin (actb). The primers used for qPCR were Kicq predesigned primers (Sigma-Aldrich).

### Analysis of rate of FA uptake

Cells were treated with a mix of 200μM non-radiolabeled palmitate (BSA conjugated at 6:1) and 0.15μCi [1-^14^C] palmitic acid (American Radiolabeled Chemicals, ARC 0172A) conjugated with 10μM BSA for 15, 30, 60 mins. Following the treatment, the cells were solubilized with RIPA lysis buffer. The ^14^C counts per min of each sample was calculated in a scintillation counter. The radioactive counts was normalized to protein concentration of the samples quantified by Bradford assay.

#### Microsome isolation

The cell pellets were dissolved in 1ml cell lysis buffer (20mM tricine and 250 mM sucrose, 1mM EDTA, pH 7.8) containing protease inihibitor cocktail (ThermoFisher, 78430). The resuspended cells were then dounce homogenized and centrifuged for 10 mins at 8000 g. The supernatant was collected and centrifuged for 1 hr at 100,000 g. The resultant microsomal pellet was resuspended in 0.1 M PK buffer (0.1 M K2HPO4 buffer set at pH 7.2 with NaH2PO4 solution).

#### SCD1 enzyme activity

The protocol was provided by Dr. James Ntambi’s lab and was also adapted from [10]. In brief, we made a reaction mix consisting of 0.03 mM stearoyl-CoA (Sigma, S0802), 1 μCi/ml radiolabelled [9,10-^3^H] stearoyl-CoA (American Radiolabeled Chemicals, ART 0390), 2 mM NADH (Sigma, N8129) in 0.1 M PK buffer (0.1 M K2HPO4 buffer set at pH 7.2 with NaH2PO4 solution). 1.5 mM of stearoyl-CoA stock was prepared in cold and fresh everytime in 10mM sodium acetate in 50% ethanol solution. The reaction assay consisted of 100 μg microsomes in 0.1 M PK buffer and 180 μl of premix. 4 μM SCDi was added as a control sample prior to the reaction. The reaction was done at 37°C for 30 mins, after which 200 μl of 6% perchloric acid was used to quench the reaction. Next 700 μl of 10% (w/v) charcoal dissolved in PBS was added to sediment the unused substrate. The sample was vigorously vortexed to mix and then centrifuged at 13000 rpm for 10 mins. The radioactive ^3^H following the SCD1 enzyme activity released in the aqueous phase which was counted in a scintillation counter. The results were quantified as ^3^H counts per min normalised to WT microsomes.

### Neutral lipid analysis by thin layer chromatography (TLC)

A protocol from [53] was used to extract lipids from cultured cells. Following treatment cells were washed twice with PBS, scraped and their pellet weights were detected. The pellets were treated with chloroform, methanol, 500 mM NaCl in 0.5% acetic acid such that the ratio of chloroform: methanol: water was 2:2:1.8. The suspended lysed cells were vortexed and centrifuged at 4000 rpm for 15 min at 4°C. The bottom chloroform layer consisting of lipids was collected, dried and then resuspended in chloroform to a volume proportional to cell pellet weight. These lipids in chloroform were separated using thin layer chromatography (TLC). The solvent used to separate neutral lipids and FFAs was hexane: diethyl ether: acetic acid (80:20:1) or cyclohexane: ethylacetate (1:2). To develop the lipid spots, the TLC plates were sprayed with 3% copper acetate dissolved in 8% phosphoric acid and heated in the oven at 135°C for ∼1hr. The spraying and heating of the TLC plates was repeated until the lipids could be properly visualized. Once the bands developed, the TLC plates were scanned and then Fiji (ImageJ) was used to quantify the intensity of the bands. The relative change in intensity of each band was quantified with respect to untreated WT sample.

### APEX2 dependent biotinylation

This protocol was modified from [39]. In brief, U2OS and HEK293 cells expressing APEX2 tagged constructs and non-APEX2 construct (negative control) were incubated with 500 μM biotin-phenol (BP) (Adipogen, CDX-B0270-M100) for 30 mins and 1mM H2O2 was added for 1 min to biotinylate proteins proximal (labeling radius of ∼20nm) to the expression of APEX2 tagged construct. The biotinylation reaction was quenched after 1 min by washing the cells thrice with a quencher solution (10mM sodium ascorbate, 5mM Trolox and 10mM sodium azide solution in DPBS). To visualize the biotinylated proteins, the cells were IF-stained with streptavidin-alexa 647 (Thermofisher Scientific, S21374) during incubation with secondary antibody and then imaged using confocal miroscopy.

To identify the biotinylated proteins, immunoprecipitation and mass spectrometry techniques were used. After the labelling reaction, the cells were scraped in quencher solution and lysed in RIPA lysis buffer consisting of protease inhibitor cocktail (ThermoFisher, 78430), 1mM PMSF and quenchers. The lysates were vortexed vigorously in cold for 30 mins and cleared by centrifuging at 10,000g for 10 mins at 4°C. The cell lysate was then dialysed using Slide-A-Lyzer dialysis cassette (3500 MWCO, ThermoFisher Scientific) to remove unreacted free BP. The protein concentration was then measured using Pierce 660nm assay. 2 mg protein was incubated with 80ul streptavidin magnetic beads (Pierce, 88817) for 1 hour at room temperature on a rotator. The beads were pelleted using a magnetic rack. The pelleted beads were washed with a series of buffers - twice with RIPA lysis buffer, once with 1 M KCl, once with 0.1 M Na_2_CO, once with 2 M urea solution in 10 mM Tris-HCl, pH 8.0, and twice with RIPA lysis buffer. To elute the biotinylated proteins, the beads were boiled in 80ul of 2X Laemmli Sample Buffer (Bio-Rad, 161-0737) supplemented with 2mM biotin and 20mM DTT. Some of the eluate was run in SDS page gel and Coomassie stained to visualize the amount of enriched protein. Some eluate was western blotted with streptavidin-HRP antibody to visualize protein biotinylation. The rest was processed for gel digestion and mass spectrometry (MS) analysis to identify the enriched biotinylated proteins.

### Immunoprecipitation (IP)

To test co-IP of SCD1 with Snx14 constructs tagged with 3XFlag at the C-terminal, the cells expressing those Snx14 constructs were lysed in IPLB buffer (20mM HEPES, pH7.4, 150mM KOAc, 2mM Mg(Ac)2, 1mM EGTA, 0.6M Sorbitol,) supplemented with 1% w/v digitonin and protease inhibitor cocktail (PIC) (ThermoFisher, 78430), followed by pulse sonication. The lysate is clarified by centrifugation at 10,000g for 10 mins. 2% of the supernatant was saved as the input. The rest of the supernatant was incubated with Flag M2 affinity gel (Sigma, A2220) for 2hrs at 4°C. The beads were then washed thrice with IPLB wash buffer (IPLB, 0.1% digitonin, PIC) and twice with IPLB without digitonin. The IPed proteins are eluted from the beads by incubating the beads with 400 μg/ml 3X Flag peptide in IPLB buffer for 30mins. The beads were spun down and the supernatant was collected. The supernatant and the input fraction was run in a SDS-page gel and western blotted for Flag constructs and SCD1.

### Western blot

Whole cell lysate was prepared by scraping cultured cells, lysing them with RIPA lysis buffer containing protease inhibitor cocktail (ThermoFisher, 78430) and pulse sonication, and then clearing by centrifugation at 10,000g for 10 mins at 4°C. Samples were prepared in 4X sample loading buffer containing 5% β-mercaptoethanol and boiled at 95^0^C for 10 mins and then run on a 10% polyacrylamide gel. Wet transfer was run to transfer the proteins from the gel onto a PVDF membrane. The proteins on the membrane were probed by blocking the membrane with 5% milk in TBS-0.1%Tween (TBST) buffer for 1 hr at RT, incubating with primary antibodies overnight, washing with TBST thrice, incubating with HRP conjugated secondary antibodies (1:5000) and developing with Clarity™ Western ECL blotting substrate (Bio-Rad, 1705061) and imaging with X-ray film. The primary antibodies used for western blot were Rab anti-EGFP (1:2000), Mouse anti-Hsp90B1(1:1000), Mouse anti-Flag(1:1000), rabbit anti-SCD1 (1:500), guinea-pig PLIN3 (Progen, GP32), streptavidin-HRP (1:1000; Thermo Scientific, S911).

For quantitative densitometry, relative protein expression was quantified by analyzing band intensities using ImageJ (Fiji) and quantifying the fold change relative to untreated WT.

### Lipidomic profiling methods

#### Lipid analysis

All solvents were either HPLC or LC/MS grade and purchased from Sigma-Aldrich. All lipid extractions were performed in 16×100mm glass tubes with PTFE-lined caps (Fisher Scientific). Glass Pasteur pipettes and solvent-resistant plasticware pipette tips (Mettler-Toledo) were used to minimize leaching of polymers and plasticizers. Fatty acid (FA) standards (FA(16:0{^2^H_31_}, FA(20:4ω6{^2^H_8_}) and FA(22:6ω3{^2^H_5_}) were purchased from Cayman Chemical, and used as internal standards for total fatty acid analysis by GC-MS. SPLASH Lipidomix (Avanti Polar Lipids, Alabaster, AL, USA) was used as internal standards for lipidomic analysis by LC-MS/MS.

#### Total Fatty Acid Analysis by GC-MS

Aliquots equivalent to 200k cells were transferred to fresh glass tubes for liquid-liquid extraction (LLE). The lipids were extracted by a three phase lipid extraction (3PLE) [54]. Briefly, 1mL of hexane, 1mL of methyl acetate, 0.75mL of acetonitrile, and 1mL of water was added to the glass tube containing the sample. The mixture was vortexed for 5 seconds and then centrifuged at 2671*×g* for 5 min, resulting in separation of three distinct liquid phases. The upper (UP) and middle (MID) organic phases layers were collected into separate glass tubes and dried under N_2_. The dried extracts were resuspended in 1mL of 0.5M potassium hydroxide solution prepared in methanol, spiked with fatty acid internal standards, and hydrolyzed at 80°C during 60 minutes. Hydrolyzed fatty acids were extracted by adding 1mL each of dichloromethane and water to the sample in hydrolysis solution. The mixture was vortexed and centrifuged at 2671*×g* for 5 minutes, and the organic phase was collected to a fresh glass tube and dried under N_2_. Total fatty acid profiles were generated by a modified GC-MS method previously described by *[55]*. Briefly, dried extracts were resuspendend in 50µL of 1% triethylamine in acetone, and derivatized with 50µL of 1% pentafluorobenzyl bromide (PFBBr) in acetone at room temperature for 25 min in capped glass tubes. Solvents were dried under N_2_, and samples were resuspended in 500µL of isooctane. Samples were analyzed using an Agilent 7890/5975C (Santa Clara, CA, USA) by electron capture negative ionization (ECNI) equipped with a DB-5MS column (40m × 0.180mm with 0.18µm film thickness) from Agilent. Hydrogen was used as carrier gas and injection port temperature were set at 300°C. Fatty acids were analyzed in selected ion monotoring (SIM) mode. The FA data was normalized to the internal standards. Fatty acid with carbon length; C ≤ 18 were normalized to FA(16:0{^2^H_31_}), C = 20 were normalized to FA(20:4 ω6{^2^H_8_}), and C = 22 were normalized to FA(22:6 ω3{^2^H_5_}). Data was processed using MassHunter software (Agilent). Abundance of lipids in each of the samples are normalized to cell number.

#### Lipidomic analysis by LC-MS/MS

Aliquots equivalent to 200k cells were transferred to fresh glass tubes for LLE. Samples were dried under N_2_ and extracted by Bligh/Dyer [56]; 1mL each of dichloromethane, methanol, and water were added to a glass tube containing the sample. The mixture was vortexed and centrifuged at 2671*×g* for 5 min, resulting in two distinct liquid phases. The organic phase was collected to a fresh glass tube, spiked with internal standards and dried under N_2_. Samples were resuspendend in Hexane.

Lipids were analyzed by LC-MS/MS using a SCIEX QTRAP 6500^+^ equipped with a Shimadzu LC-30AD (Kyoto, Japan) HPLC system and a 150 × 2.1 mm, 5µm Supelco Ascentis silica column (Bellefonte, PA, USA). Samples were injected at a flow rate of 0.3 ml/min at 2.5% solvent B (methyl tert-butyl ether) and 97.5% Solvent A (hexane). Solvent B is increased to 5% during 3 minutes and then to 60% over 6 minutes. Solvent B is decreased to 0% during 30 seconds while Solvent C (90:10 (v/v) Isopropanol-water) is set at 20% and increased to 40% during the following 11 minutes. Solvent C is increased to 44% for 6 minutes and then to 60% during 50 seconds. The system was held at 60% of solvent C during 1 minutes prior to re-equilibration at 2.5% of solvent B for 5 minutes at a 1.2mL/min flow rate. Solvent D (95:5 (v/v) Acetonitrile-water with 10mM Ammonium acetate) was infused post-column at 0.03ml/min. Column oven temperature was 25°C. Data was acquired in positive and negative ionization mode using multiple reaction monitoring (MRM). Each lipid class was normalized to its correspondent internal standard. The LC-MS data was analyzed using MultiQuant software (SCIEX). Abundance of lipids in each of the samples are normalized to cell number.

## Acknowledgements

The authors would like to thank Joel Goodman, Sandra Schmid, Russell Debose-Boyd, and members of the Henne lab for help and conceptual advise in the completion of this study. We would also like to thank Dr. Philip Stanier and Dr. Dale Bryant (UCL, London, UK) for providing the SCAR20 patient cells. We acknowledge Dr. Kate Luby-Phelps and Anza Darehshouri for technical assistance with confocal and electron microscopy. We thank Dr. Andrew Lemoff for assistance with mass-spectrometry proteomics. W.M.H. is supported by funds from the Welch Foundation (I-1873), the Searle Foundation (SSP-2016-1482), the NIH NIGMS (GM119768), and the UT Southwestern Endowed Scholars Program. S.D. is supported by an AHA Pre-doctoral Fellowship Grant (20PRE35210230). JGM is supported in part by NIH 2 PO1 HL20948-33 grants.

## Figure legends

**Supplementary Figure 2.**
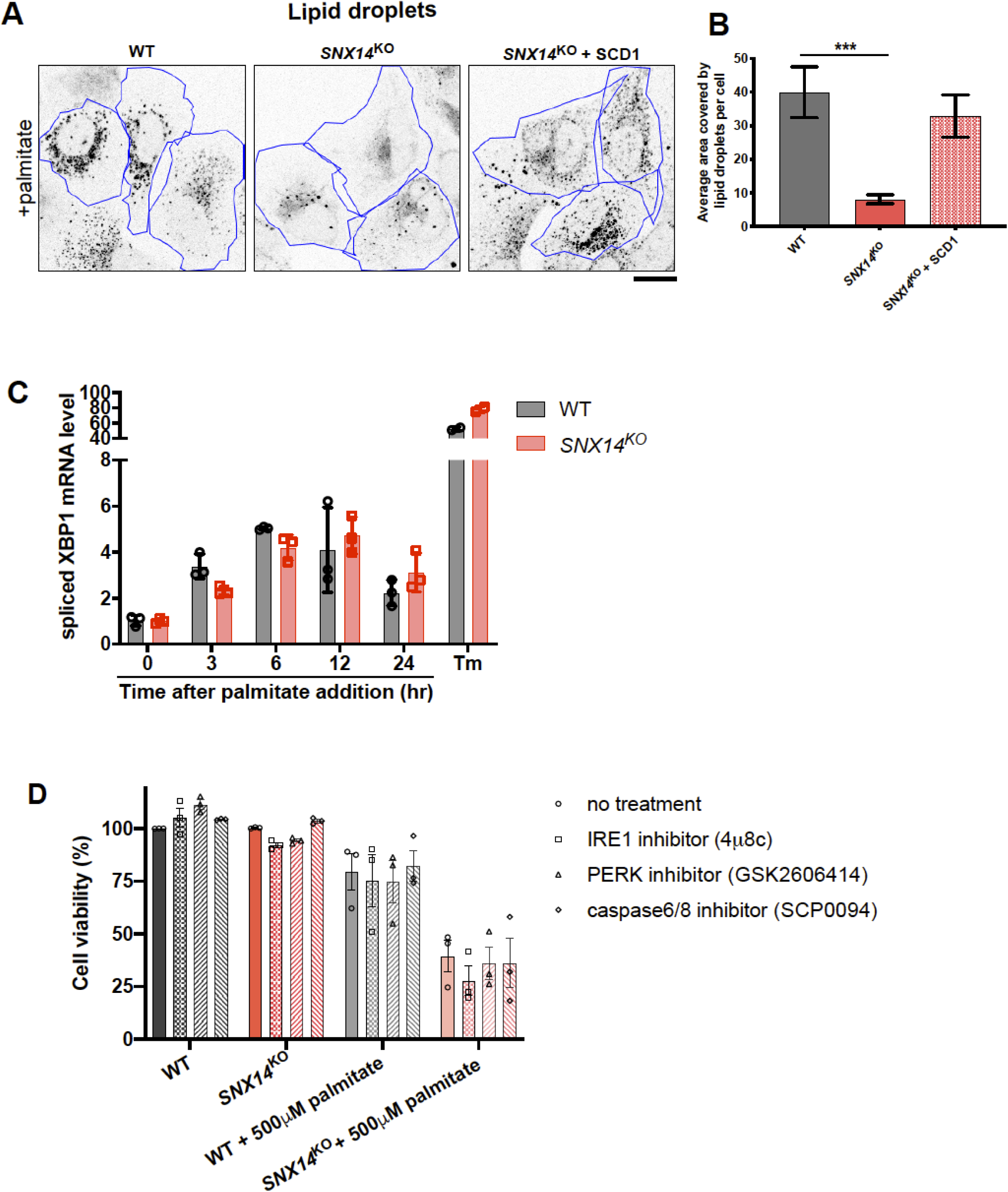
**A**. Confocal micrographs of WT and *SNX14*^KO^ cells treated with palmitate overnight and stained with monodansylpentane (MDH) to visualize LDs (black). The cell boundary is marked by blue outline. Images are processed so that the LDs are grayscaled and then inverted in Image-J. Scale bar = 10μm. **B**. Average area covered by LDs per cell of representative images from Fig S2A is quantified. Total LD area was derived from six fields of view, each consisting of approximately eight cells from two different sets of experiments (total no. of cells ∼90; ***p < 0.0001 unpaired t test with α = 0.05). **C**. RT-PCR data of spliced Xbp1 mRNA levels which is an indicator of UPR activity in WT and *SNX14*^KO^ cells treated with palmitate for indicated time. Tunicamycin (Tm) treatment for 6hr (5 µg/ml) used as control which induces UPR activity. **D**. Cell viability (%) of WT and *SNX14*^KO^ cells following 2 days exposure to 500μM palmitate and either untreated or treated with one of the following –a) 64μM of IRE1 inhibitor 4μ8c (b) 30 nM of PERK inhibitor GSK2606414 (c) 40μM of caspase 6/8 inhibitor SCP0094. The assay was repeated thrice in triplicates. Values represent mean±SEM.

**Supplementary Figure 3.**
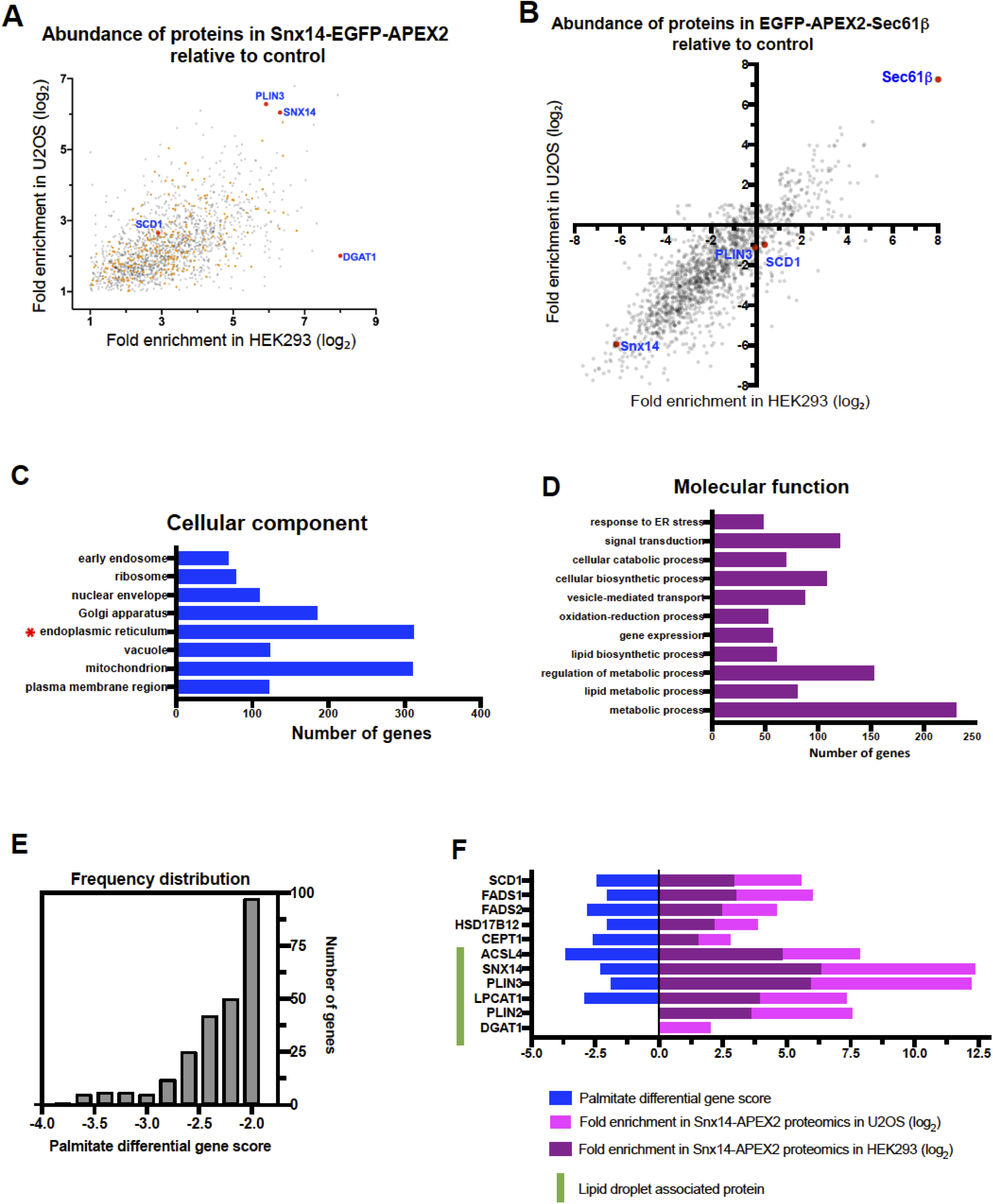
**A**. Proteins enriched for Snx14 from set (1) of Fig 3E after excluding proteins also enriched for Sec61 are plotted as log_2_ fold enrichment relative to negative control from HEK293 (x-axis) and U2OS cells (y-axis) and are denoted by grey dots. Orange dots represent those proteins whose CRISPR/Cas9 deletion results in palmitate mediated sensitivity. **B**. Log_2_ fold enrichment of proteins in APEX2-Sec61 relative to negative control from HEK293 (x-axis) and U2OS cells (y-axis) are plotted and denoted by grey dots. **C**. Gene ontology enrichment analysis by ClueGO using Cytoscape software grouped genes from set (1) of Fig 3E according to their association with a cellular organelle. **D**. Gene ontology enrichment analysis by ClueGO using Cytoscape software clustered ER-localized genes from Fig S3C according to their molecular function. **E**. Frequency distribution of the CRISPR/Cas9 screened genes tested negative (<-1.8) for palmitate treatment from [46] enriched specifically for Snx14 (denoted as orange dots) in Fig S3A. **F**. Examples of some proteins including LD-associated proteins which are specifically and highly enriched in Snx14 after analysis in Fig 3E and also has a high negative screen score showing palmitate sensitivity similar to Snx14 from [46].

**Supplementary Figure 4.**
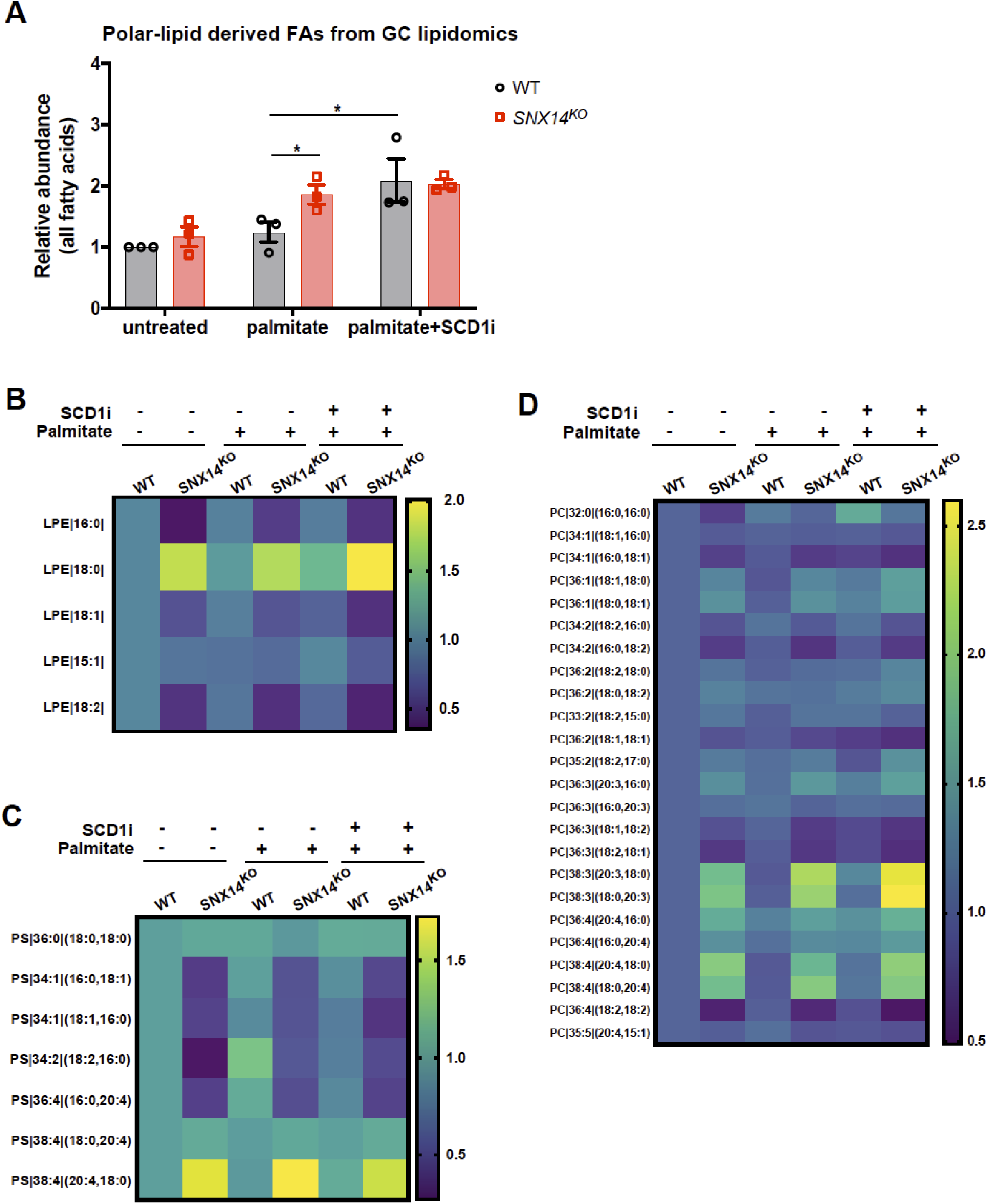
**A**. Abundance of all fatty acids derived from polar lipids of WT and *SNX14*^KO^ cells relative to untreated WT under the following conditions – no treatment, palmitate treatment or treatment with palmitate and SCD1i. Values represent mean±SEM (n=3, *p<0.01, multiple t-test by Holm-Sidak method with alpha = 0.05). B. Heatmap indicating the relative change in abundance of 5 different LPE species of WT and *SNX14*^KO^ cells relative to untreated WT. These cells were either untreated or treated with palmitate or palmitate in presence of SCD1i. C. Heatmap indicating the relative change in abundance of 7 different PS species of WT and *SNX14*^KO^ cells relative to untreated WT. These cells were either untreated or treated with palmitate or palmitate in presence of SCD1i. D. Heatmap indicating the relative change in abundance of 24 different PC species of WT and *SNX14*^KO^ cells relative to untreated WT. These cells were either untreated or treated with palmitate or palmitate in presence of SCD1i.

**Supplementary Figure 5.**
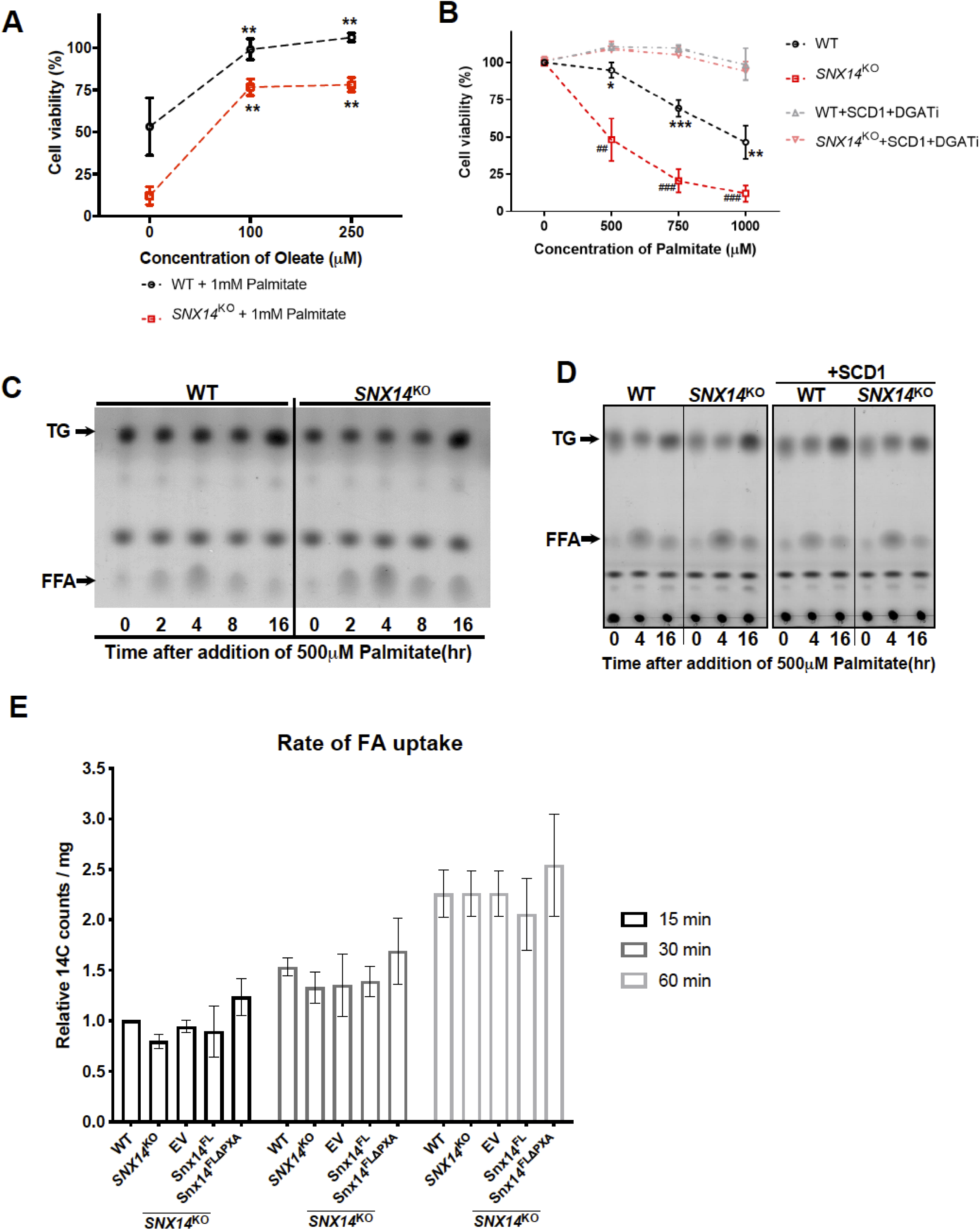
A. Cell viability (%) shows increase in surviving WT and *SNX14*^KO^ cells when treatment with 1mM palmitate is supplemented with 100 and 200μM of oleate. The assay was repeated thrice in triplicates. Values represent mean±SEM (**p<0.001, multiple t-test by Holm-Sidak method with alpha = 0.05, significance test between oleate treatment with non-oleate treatment). B. Cell viability (%) of WT and *SNX14*^KO^ cells, showing sensitivity of both WT and *SNX14*^KO^ cells are reduced with reintroduction of SCD1 in presence of DGAT1/2 inhibitors (DGATi) following addition of increasing concentration (0, 500, 750, 1000μM) of palmitate for 2 days. The assay was repeated thrice in triplicates. Values represent mean±SEM. Significance test between WT and WT+SCD1+DGATi denoted as * (n=3, *p<0.01, **p<0.001, ***p<0.0001, multiple t-test by Holm-Sidak method with alpha = 0.05) Significance test between *SNX14*^KO^ and *SNX14*^KO^+SCD1+DGATi denoted as ^#^ (n=3, ^##^p<0.001, ^##^ p<0.0001, multiple t-test by Holm-Sidak method with alpha = 0.05). **C**. TLC of neutral lipids and FFAs performed in WT and *SNX14*^KO^ cells treated with 500μM palmitate for 0, 2, 4, 8, 16 hours. **D**. TLC of neutral lipids and FFAs performed in WT and *SNX14*^KO^ cells before and after overexpression of SCD1 following exposure to 500μM palmitate for 0, 4, 16 hours. **E**. Fatty acid uptake in WT, *SNX14*^KO^ cells and *SNX14*^KO^ cells expressing either EV, Snx14^FL^, or Snx14^FLΔPXA^ following exposure to ^14^C palmitate for 15, 30, 60 mins quantified relative to WT treated with 15 min ^14^C palmitate.

**Supplementary Figure 6.**
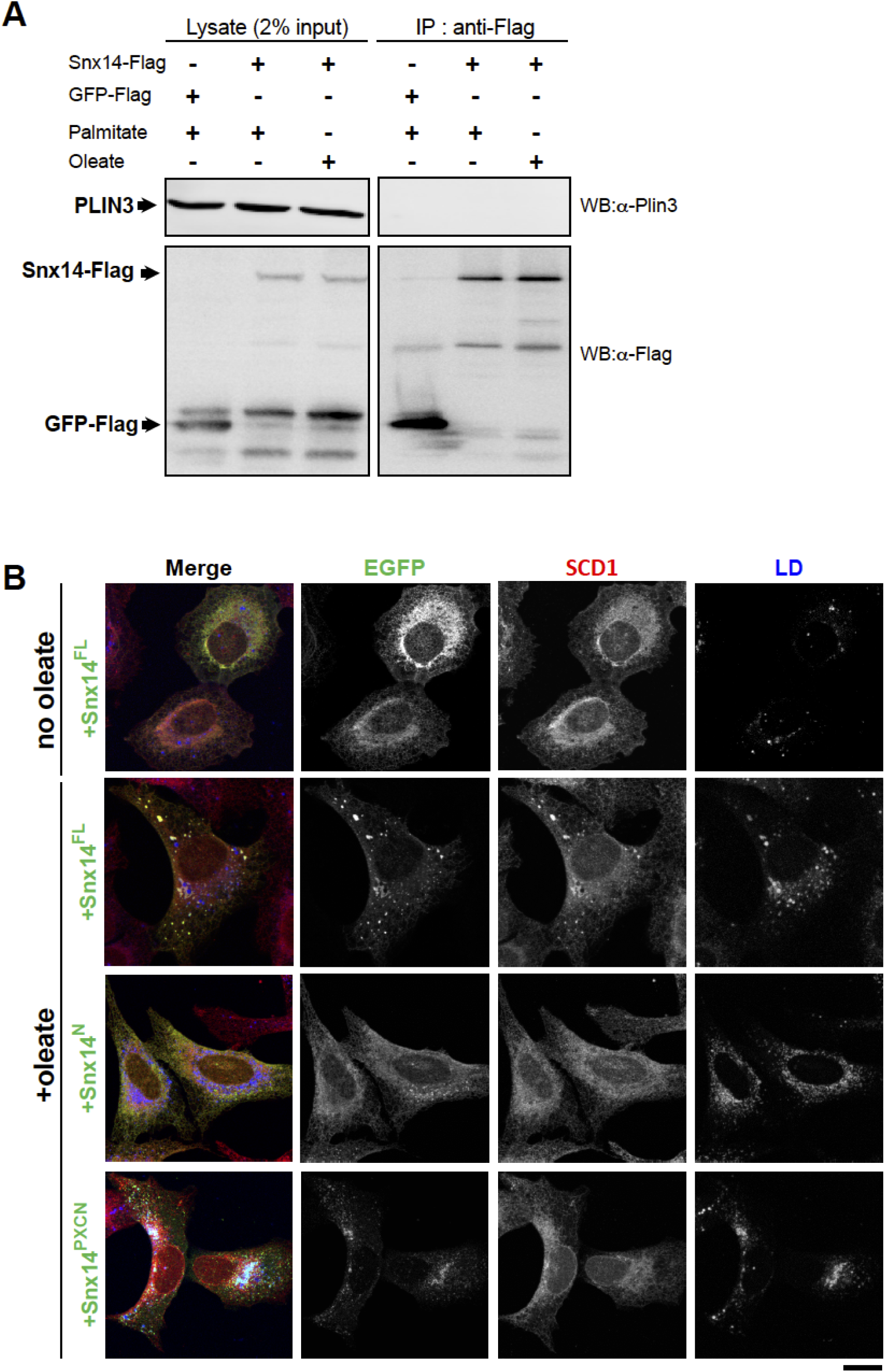
**A**. Lanes represent 2% input and IP from GFP-Flag and Snx14^FL^-3XFlag expressing U2OS cells with oleate and palmitate treatment respectively. Western blotting with anti-Flag and anti-PLIN3 antibody reveals relative expression of all the Flag tagged constructs and PLIN3 in all these samples. Snx14^FL^-3XFlag similar to GFP-Flag could not co-IP PLIN3. **B**. IF staining of U2OS cells expressing Snx14^FL^, Snx14^N^, Snx14^PXCN^ with anti-EGFP(green), anti-SCD1(red) antibody and imaged with confocal microscope. LDs were stained with MDH (blue). The cells were either untreated or treated with oleate. Scale bar = 10 μm.

